# Differential impact of Kv8.2 loss on rod and cone signaling and degeneration

**DOI:** 10.1101/2021.07.05.451197

**Authors:** Shivangi M Inamdar, Colten K Lankford, Deepak Poria, Joseph G Laird, Eduardo Solessio, Vladimir J. Kefalov, Sheila A Baker

## Abstract

Heteromeric Kv2.1/Kv8.2 channels are voltage-gated potassium channels localized to the photoreceptor inner segment. They carry *I*_Kx_, which is largely responsible for setting the photoreceptor resting membrane potential. Mutations in Kv8.2 result in childhood-onset Cone Dystrophy with Supernormal Rod Response (CDSRR). We generated a Kv8.2 knockout (KO) mouse and examined retinal signaling and photoreceptor degeneration to gain deeper insight into the complex phenotypes of this disease. Using electroretinograms we show that there were delayed or reduced signaling from rods depending on the intensity of the light stimulus, consistent with reduced capacity for light-evoked changes in membrane potential. The delayed response was not seen *ex vivo* where extracellular potassium levels were controlled by the perfusion buffer, so we propose the *in vivo* alteration is influenced by genotype-associated ionic imbalance. We observed mild retinal degeneration. Signaling from cones was reduced but there was no loss of cone density. Loss of Kv8.2 altered responses to flickering light with responses attenuated at high frequencies and altered in shape at low frequencies. The Kv8.2 KO line on an all-cone retina background had reduced cone-driven ERG b wave amplitudes and underwent degeneration. Altogether, we provide insight into how a deficit in the dark current affects the health and function of photoreceptors.

## INTRODUCTION

The Kv2 subfamily of voltage-gated K^+^ channels are widely expressed. In general, they carry a delayed rectifier current but incorporation of a regulatory subunit, known as an electrically silent (KvS) subunit can dramatically shift the activity profile of Kv2 containing channels (1). There are 10 KvS subunits that differ by tissue expression. To date, only one is known to cause disease. The KvS subunit, Kv8.2, is expressed in photoreceptors and mutations in the gene encoding Kv8.2, *KCNV2*, are the cause of Cone Dystrophy with Supernormal Rod Responses (CDSRR) (2,3). CDSRR is also known as KCNV2 retinopathy or Retinal Cone Dystrophy 3B (OMIM # 610356).

CDSRR is fascinating because photoreceptor signaling is perturbed differently as a function of light intensity. Using electroretinography (ERG), which is essential to diagnose this disease, different changes in the b wave amplitude are seen. In very dim light rods underperform, while brighter light triggers a ‘supernormal rod response’. Cone-driven responses are reduced throughout their functional range (2). The disease is typically diagnosed in school age children. Common symptoms include loss of visual acuity and color blindness, photophobia, and progressive macular atrophy (3–8). A significant patient population also suffers from night blindness further distinguishing this disease from other types of cone dystrophies (4).

Only recently have the first animal models for the study of CDSRR been described. Kv8.2 assembles with Kv2.1 in photoreceptors to form a hetero-tetrameric channel that carries *I*_Kx_ (3,9,10). In studies of Kv8.2 knockout (KO) mice at young or mid-age, ERG analysis was used to show that these mice phenocopy human CDSRR. While mice don’t have a macula, both rod and cone degeneration were reported (11,12).

An elegant combination of electrophysiological recordings and biophysical modeling using the Kv2.1 KO mouse provided direct evidence that Kv2.1-containing channels are necessary for the outward component of the circulating dark current. i.e. *I*_Kx_ (13). With the loss of Kv2.1, rod responses were reduced. The remaining fraction of current is supported primarily by the activity of the electrogenic Na^+^/K^+^-ATPase pump. Moreover, the resting membrane potential in Kv2.1 KO rods was depolarized. This condition allows excessive calcium to enter outer segments through CNG channels, thus explaining the mitochondrial dysgenesis and slow retinal degeneration occurring in these mice.

Kv8.2 trafficking to the plasma membrane requires Kv2.1 (14). Therefore, in the Kv2.1 KO, Kv8.2 should not be available at the surface to influence rod signaling. This may mean that Kv2.1 KO mice would also have a supernormal b wave, but that has been reported in only two of the three studies (11–13). Kv2.1 trafficking is independent of Kv8.2 so loss of Kv8.2 should result in photoreceptors expressing homomeric Kv2.1 channels, that are largely inactive in the normal operational range of photoreceptors. However, if loss of Kv8.2 depolarizes the resting membrane potential then loss of Kv8.2 function could lead to an altered current from homomeric Kv2.1 channels. The uncertainties regarding rod and cone signaling upon loss of Kv8.2 is compounded by the absence of any information regarding how macular degeneration is triggered.

We investigated retinal signaling, defined by the summed electrical activity of the retina as measured by ERG, in an independent Kv8.2 KO mouse. We confirm the overall ERG phenotype as reported in the complementary Kv8.2 KO mouse line (11,12). We obtained different results for two experiments – we observed an increase in Kv2.1 expression and there was no loss of cone density in our mouse line. We extended the understanding of this model for CDSRR by using *ex vivo* ERG to measure isolated photoreceptor responses, by investigating responses to flickering light stimuli, and by testing for photophobia. We also crossed the Kv8.2 KO to a mouse line having an all-cone retina where we observed reduced cone-driven ERG responses and cone degeneration. Our work demonstrates that Kv8.2 KO photoreceptor responses are delayed and altered in amplitude. Unlike CDSSR patients, the mice lack photophobia and while rods degenerate, cones only do so when in an all-cone environment.

## RESULTS

### Generation of the Kv8.2 KO

CRISPR/Cas9 genome editing using two guides targeting the 5’ end of mouse *Kcnv2* exon 1 was performed, and two founder lines were obtained that carried the desired frameshifting 101 bp deletion as verified by sequencing (Fig. 1A). Offspring from both founder lines generated indistinguishable data so for simplicity, we used the N2 generation from the 3564.1 founder line for most of the experiments presented here. Incorporation of frameshifting mutations can result in loss of mRNA through non-sense mediated decay. To test for this, retinal RNA was reverse transcribed and transcript levels of *Kcnv2* were measured with dd-PCR. In the Kv8.2 KO retina, *Kcnv2* transcripts were reduced by 50% compared to WT littermate controls (Supplemental Fig. 1). Therefore, the genome editing altered, but did not prevent, expression of *Kcnv2* in the mutant mice.

**Figure 1:**
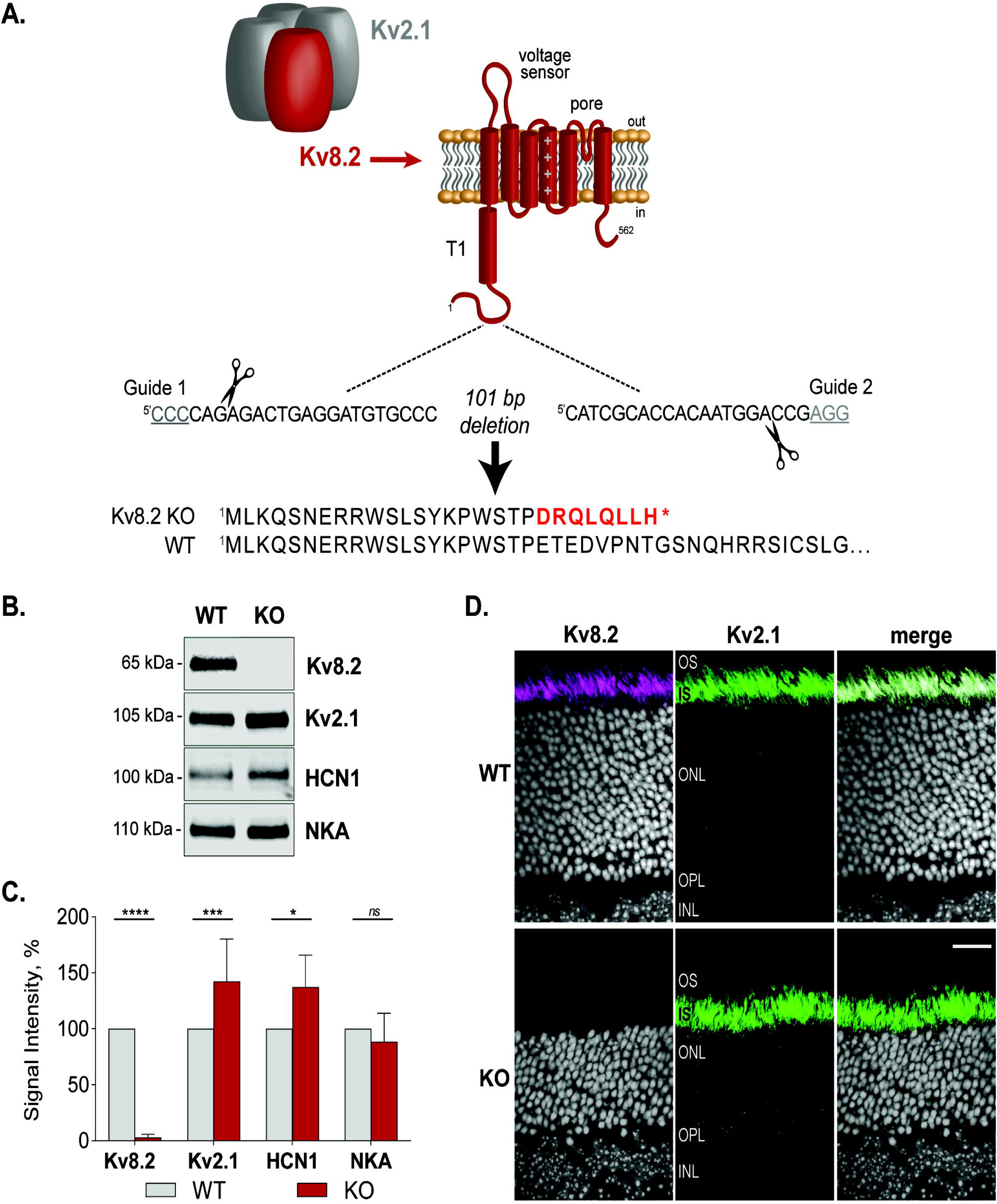
Design of the Kv8.2 KO strain. ***A)*** Kv8.2 (red) requires assembly with Kv2.1 (gray) to form a functional channel. Kv8.2 has the typical topology of a voltage-gated K+ channel subunit; a T1 tetramerization domain, voltage sensor, and pore forming domain. Guide RNAs were designed to excise 101 nt from *Kcnv2* to generate a frameshifting truncation in the N-terminus of Kv8.2 prior to any functional domains. ***B)*** Representative Western blots of retina lysates from WT vs KO animals, probed for Kv8.2, Kv2.1, HCN1, and Na/K-ATPase (NKA). ***C)*** Expression levels of proteins as measured in (*B*) normalized to WT levels. ***D)*** Immunostaining for Kv8.2 (magenta) and Kv2.1 (green) in WT versus Kv8.2 KO photoreceptors. Abbreviations are OS, outer segment; IS, inner segment; ONL, outer nuclear layer; OPL, outer plexiform layer; INL, inner nuclear layer; and the scale bar is 20 µm.

Western blotting for Kv8.2 in retina lysates confirmed the absence of protein expression (Δ mean = -97, 95% CI [-68, -126], 2-way ANOVA, adj.p < 0.0001). Expression of Kv2.1, the interaction partner of Kv8.2 was upregulated in the Kv8.2 KO retina (Δmean = 42, 95% CI [68, 16], adj.2-way ANOVA p = 0.0008). This could reflect an increase in Kv2.1 homomeric channels in compensation for the lost Kv2.1/Kv8.2 heteromeric channels. HCN1, an ion channel with a complementary function to Kv8.2 was also upregulated in the mutant mice (Δ diff = 37, 95% CI [68, 5,], 2-way ANOVA adj. p = 0.0164), but Na/K-ATPase (NKA), which contributes to resting membrane potential was unaltered (Δ diff = 11, 95% CI [-32, 56], 2-way ANOVA adj. p= 0.9290) (Fig. 1B, C).

We found the Kv8.2 antibody (Clone N448/88 from NeuroMab) generated a non-specific signal in immunohistochemistry (Supplemental Fig. 2). Courtesy of James Trimmer, we were able to screen additional clones from the original N448 monoclonal antibody project and identified clone N448/50 as specific in both Western blots and immunohistochemistry experiments. Kv8.2 and Kv2.1 are localized to the apical end of photoreceptor inner segments. In the Kv8.2 KO, the Kv8.2 signal was lost but the Kv2.1 signal was maintained (Fig. 1D). The relative thinness of the outer nuclear layer in the KO at 3.5 months of age indicates rod degeneration is occurring, which is described in greater detail in a subsequent section. Altogether, these data validate this Kv8.2 KO strain.

### Aberrant retinal activity in Kv8.2 KO mice phenocopies CDSRR patients

Electroretinograms (ERG) are used to diagnose CDSRR patients following the standardized protocol described by the International Society for Clinical Electrophysiology of Vision (ISCEV) (15). We applied this protocol to Kv8.2 KO and WT littermate controls; the results were consistent with a diagnosis of CDSRR (Fig. 2). The dark-adapted WT and KO animals did not generate significantly different responses to the dimmest flash (Δ mean ± SEM = -2.3 ± 1.4, 95% CI [-5, 0.7], t-test, p = 0.1249)) (Fig. 2A). We found that plotting the ratio of the b/a wave amplitude was the most informative. Quantitation of the individual a and b wave amplitudes for the steps where we show the ratio in Fig 2 are provided in Supplemental Fig 3. A brighter flash revealed three abnormalities in the KO response: 1) a reduced and ‘squared off’ a wave, 2) an increased b/a wave amplitude (Δ mean ± SEM = 1.6 ± 0.1, 95% CI [1, 2], t-test, p <0.0001), and 3) reduced oscillatory potentials (Δmean ± SEM = -194 ± 32, 95% CI [-264, -125], t-test, p <0.0001) (Fig. 2A-C). The contribution of cone signaling to the ERG is detected after light-adaptation in the standard combined response or the 30 Hz flicker. In both tests, the amplitude of the response was significantly reduced in KO animals ((standard combined response Δ mean ± SEM = -4.6 ± 0.53, 95% CI [-5, -3], t-test, p <0.0001), (30 Hz flicker Δ mean ± SEM = -17 ± 2, 95% CI [-22, -11], t-test, p <0.0001); (Fig. 2D, E)).

**Figure 2:**
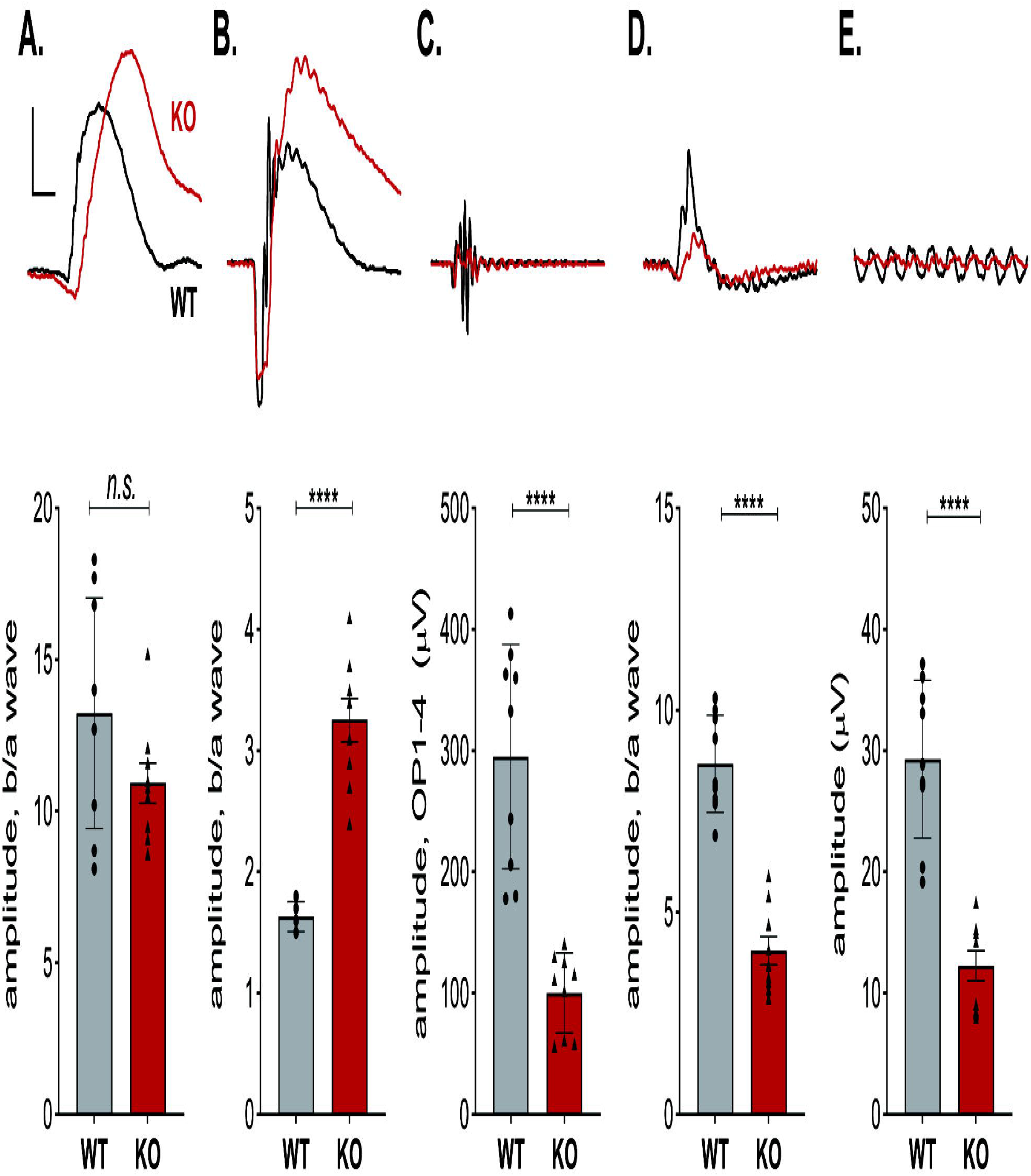
Retinal signaling in Kv8.2 KO animals. ERG responses following the standard ISCEV protocol were collected from 9 WT (black) and 9 Kv8.2 KO (red) animals ranging in age from 1-4 months. Representative traces are shown above quantitation of the amplitude of the b wave as a fraction of the a wave amplitude for (*A-D*), or the amplitude between the peaks of the first negative and first positive deflection for (*E*). ***A)*** Scotopic dim flash, ***B)*** Scotopic bright flash, **C)** oscillatory potentials extracted from (*B*), ***D)*** standard combined response (rod plus cone), ***E)*** 30 Hz flicker. Scale is 200 µV (*A-C*) or 100 µV (*D, E*) by 40 ms.

To further explore the Kv8.2 KO dependent changes in ERG phenotype we recorded responses to a series of increasingly bright flashes. To investigate rod-driven responses we used dark-adapted animals (Supplemental Fig. 4). For descriptive statistics of this experiment, see Supplemental Tables S4B to S4G. The amplitude of the scotopic a wave from KO animals was slightly reduced only at the highest flash intensity (Supplemental Fig. 4B). The time to onset for the a wave was delayed at low light intensities but became normal as the intensity of the light increased (Supplemental Fig 4C). The amplitude of the b wave did not appear different however, when we normalized the b wave to the a wave, it could be seen that the KO animals had a subnormal response at the lowest light intensities, was indistinguishable from WT at -1 log (cd·s/m^2^), then switched to the characteristic supernormal response as the light intensity increased (Supplemental Fig 4D, F). The time to onset for the b wave was delayed at all tested light intensities even after subtracting out delays from the a wave (Supplemental Fig 4B, G).

To investigate in more depth cone-driven responses we used light-adapted animals (Supplemental Fig. 5). For descriptive statistics of this experiment, see Supplemental Tables S5B to S5G. Overall, the wave forms were not as different as they were for the scotopic responses (Supplemental Fig. 5A, compare to Supplemental Fig. 4A). Quantitation of the a wave shows that in Kv8.2 KO mice, the amplitude was only reduced at the highest flash intensities (Supplemental Fig. 5B). Note, however, that the signal to noise ratio was low at the dimmer test flashes making it difficult to analyze the response. The b wave amplitudes were reduced at all intensities (Supplemental Fig 5D). When we analyzed the b/a ratio, these differences were not statistically significant (Supplemental Fig. 5F). This indicates that the changes in the cone-driven b wave were caused by reductions in the a wave. The time of onset for the a wave appeared delayed in the Kv8.2 KO (Supplemental Fig. 5C) but this was only statistically significant at two test flashes and given the small amplitude of the a wave and the potential technical error in measurement, this may not be of biological significance. The delay in the onset of the b wave was more pronounced and statistically significant at the brighter test flashes, including after subtracting out delays in the onset of the a wave (Supplemental Fig. 5E, G).

### Temporal properties of signaling in the Kv8.2 KO retina

A feature of both rod- and cone-driven signaling as documented above is a delay in response time. This is consistent with the role that *I*_kx_ plays in modulating the temporal properties of photoreceptors by accelerating the voltage response particularly at the low end of the photoreceptor dynamic range. To further probe the requirement for Kv8.2 in the temporal properties of the retina, we recorded ERG using a flickering light stimulus.

In the ISCEV protocol the Kv8.2 KO mice had reduced responses to a repetitive (0.5 log (cd.s/m^2^)) flash at 5 Hz (data not shown) and 30 Hz (see Fig. 2E). To further examine the dynamics at additional frequencies, dark-adapted mice were subjected to repetitive (0.5 log (cd.s/m^2^)) flashes of light with frequencies ranging from 0.5 Hz to 30 Hz. (Supplemental Figure 6 with descriptive statistics in Supplemental Table S6). At 0.5 Hz, the KO response was similar to that obtained with a single flash under dark adapted conditions: a reduced and squared-off a wave with a supernormal b wave. Above 0.5 Hz, the KO responses were reduced in a frequency-dependent manner with KO responses being 70% that of controls at 1 Hz, roughly 50% from 2-20 Hz, and 30% at 30 Hz, consistent with the results shown in Fig. 2.

In this standard flicker protocol, the temporal signaling properties of the Kv8.2 KO appear compromised. However, as the frequency increases there is also an effective increase in average background light. To address this issue, we used an alternate flicker ERG protocol where the stimulus is sinusoidally modulated so that the background light intensity remains constant throughout the frequency range being tested (16,17). We set the background light to mesopic levels (log (0 cd/m^2^)) and the frequency range from 0.5 to 25 Hz. This resulted in a complex waveform at low frequencies which became more sinusoidal as the frequency increased with WT and KO generating distinctly different patterns at low frequencies (Fig. 3A). For descriptive statistics of this experiment, see Supplemental Tables S3B, S3D, S3E). We determined the fundamental component derived with a Fourier transform and plotted the amplitude of the fundamental (f_0_) as a function of stimulus frequency (Fig. 3B). The fundamental amplitude was highest at 0.5 Hz, with a secondary peak at 9 Hz and reduced sharply at frequencies > 9 Hz in both WT and Kv8.2 KO. Kv8.2 KO response amplitudes were significantly reduced at all frequencies.

**Figure 3:**
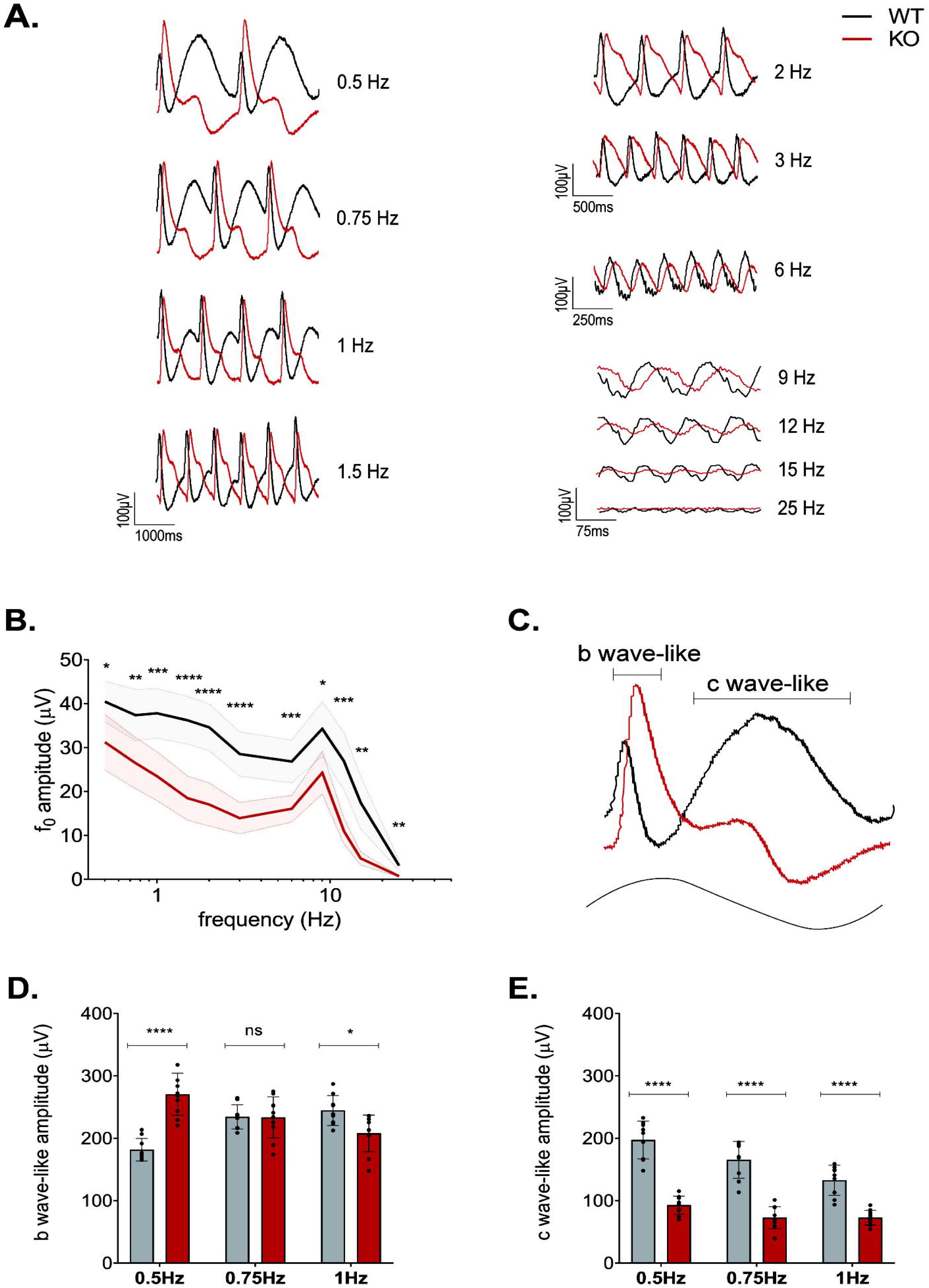
Flicker ERG reveals complex alteration to dynamic signaling in Kv8.2 KO. ***A)*** Representative ERG traces of WT (black) and Kv8.2 KO (red) to a sinusoidal flicker of increasing frequency at log 0 cd/m^2^ with 100% contrast. ***B)*** Amplitude of the fundamental component (f_0_) as a function of the flicker frequency. ***C)*** Single cycle response to a 0.5 Hz flicker with the ‘b wave-like’ and ‘c wave-like’ components of the response shown with brackets. ***D)*** Amplitude of the b wave-like component and ***(E)*** c wave-like component at 0.5, 0.75 and 1 Hz.

Analysis using the fundamental magnitude may be limited by the non-linearity of the low frequency responses as well as the distinctly different response shapes elicited by the Kv8.2 KO and control mice. We noted that the low frequency (0.5-1 Hz) responses appeared to be composed of two primary positive-going components, a sharp peak and a broad peak. The sharp peak originates with the rising phase of the sinusoidal stimulus while the broad peak occurs with decrement of the sinusoidal stimulus. These components resemble the b wave and c wave of a response to a single flash and henceforth we refer to them as “b wave-like” (sharp peak) and “c wave-like” (broad peak) (Fig. 3C). Both the b wave-like and c wave-like components differed between KCNV2 KO and controls at these frequencies. The b wave-like component was elevated in the KO at 0.5 Hz, similar to the supernormal b wave observed in the 0.5 Hz, 0.5 log (cd.s/m^2^) flash flicker. But this component was identical between KO and WT at 0.75 Hz, and slightly reduced at 1 Hz (Fig. 3D). The c wave-like component was greatly reduced in the Kv8.2 KO at all three low frequencies tested (Fig. 3E) and may reflect the absence of a true c wave in the absence of Kv8.2 as measured in a separate Kv8.2 KO line as well as the Kv2.1 KO (11–13). Thus, Kv8.2 KO appears to attenuate the response amplitude to high frequency stimulation but has a complex effect on the low frequency response by altering the response waveform.

### Isolated photoreceptor responses

Reductions in the ERG a wave amplitude may be a direct effect of the loss of Kv2.1/Kv8.2 channels or an indirect effect from photoreceptor degeneration. To isolate photoreceptor responses, we turned to *ex-vivo* transretinal ERG which allowed us to pharmacologically block synaptic transmission (Fig. 4, Table 1). We observed that the amplitude of dark-adapted rod responses was reduced in Kv8.2 KO retinas by 34% (Fig. 4B) compared to controls (Fig. 4A; see also Fig. 4C), consistent with *in vivo* ERG findings. Normalization of the response curves revealed that the sensitivity of the rods was slightly reduced in KO retina (Fig. 4C, inset). There was no delay in the time to peak but the recovery of the response in KO retina was 28% faster. The loss of the delayed onset in the *ex vivo* ERG where KO and control retinas are perfused in the same buffer indicate that the delayed signaling observed *in vivo* is due to altered extracellular K^+^ levels in the subretinal space of the intact eyeball.

**Figure 4:**
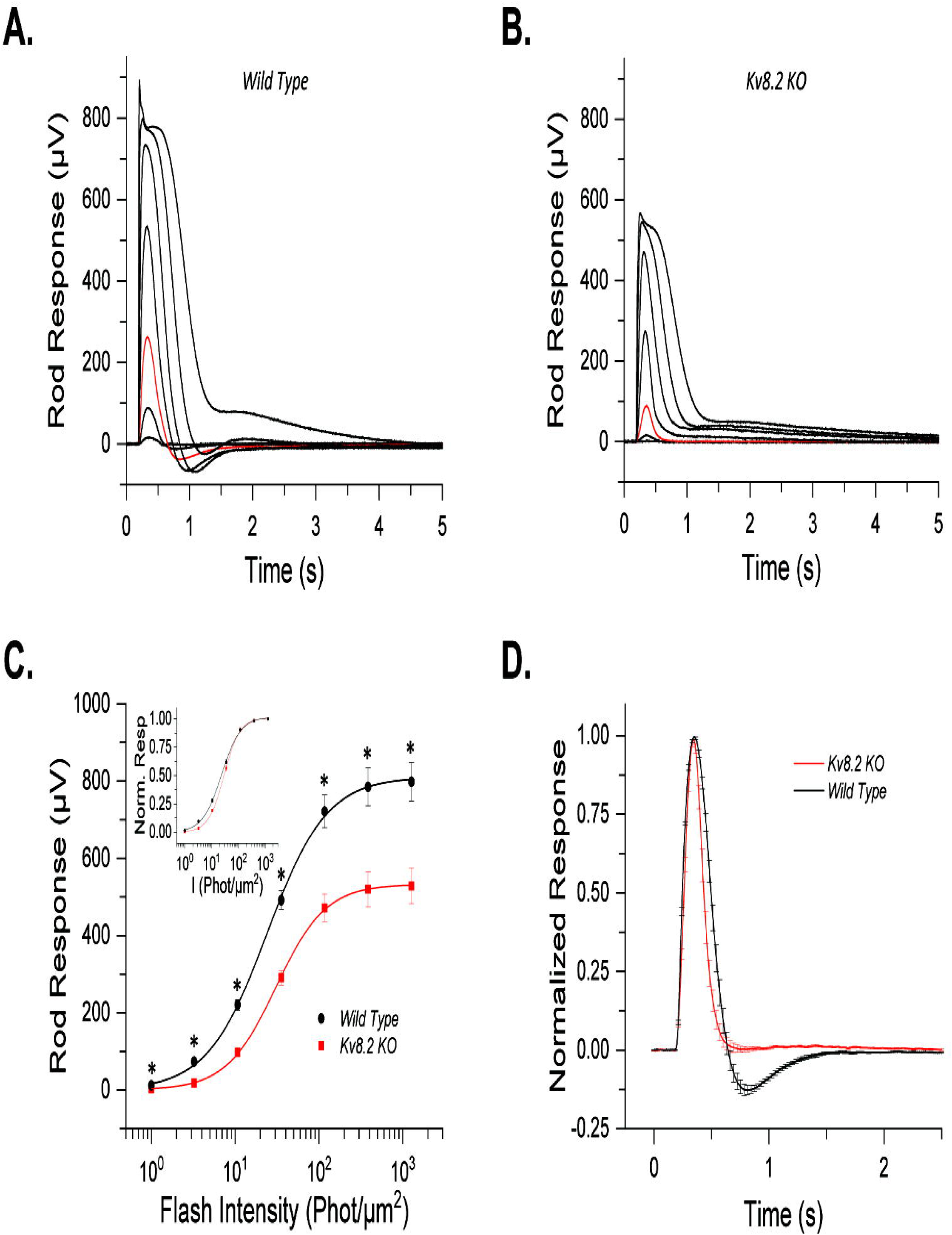
Ex-vivo responses of Kv8.2 KO rods. ***A)*** Representative responses to a family of flashes (1, 3.25, 10.7, 35.5, 117, 386 and 1271.5 photons µm^-2^) from wild-type control and ***B)*** Kv8.2 KO rods. The red traces in each case represent the dim flash response to 10.7 photons µm^-2^, highlighted for comparison. ***C)*** Averaged intensity response curves for Kv8.2 KO (red) and control (black) rods fit to a Naka-Rushton function; ***inset)*** normalized intensity response curves. ***D)*** Normalized dim flash responses from the Kv8.2 KO rods and the wild-type rods plotted together for comparison.

**Table 1:** Rod response parameters from ex vivo recordings.

| | $R_{max}$ (μV) | $I_{1/2}$<br>(Phot/μm<sup>2</sup>) | $S_{\mathcal{D}}$<br>(Phot<sup>-1</sup>/μm<sup>-2</sup>) | $t_p$ (ms) | $t_{int}$ (ms) | $\tau_{rec}$ (ms) | N |
| --- | --- | --- | --- | --- | --- | --- | --- |
| WT | 798 ± 49 | 24.0 ± 1.4 | 2.6 ± 1.4 x 10<sup>2</sup> | 170 ± 4 | 452 ± 18 | 100 ± 6 | 7 retinas (5 mice) |
| Kv8.2 KO | 528 ± 46 | 30.6 ± 2.0 | 1.8 ± 0.1 x 10<sup>2</sup> | 165 ± 4 | 466 ± 40 | 72 ± 7 | 10 retinas (6 mice) |
| *P*<br>*value* | 0.0013 | 0.02 | 0.007 | 0.33 | 0.76 | 0.009 |  |

$R_{max}$ , saturated response amplitude measured at the plateau; $I_{1/2}$ , intensity required to produce half of the saturated response; $S_{\mathcal{D}}$ , dark adapted sensitivity; $t_p$ , time to peak of a dim flash response; $t_{int}$ , integration time of the response; $\tau_{rec}$ , recovery time constant during response shut off.

Progressive macular degeneration and reduced cone function is a defining feature of CDSRR patients. We also used *ex-vivo* transretinal ERG to determine if loss of Kv8.2 causes an intrinsic change in cone responses. Saturated rod responses were subtracted from test flashes to obtain cone responses in KO retinas with pharmacological blockade of synaptic transmission (Fig. 5, Table 2). The higher signal-to-noise ratio and removal of the b-wave allowed us to obtain robust photopic a wave responses from control and Kv8.2 KO retinas. We found that the amplitude of cone responses was reduced by 20% in Kv8.2 KO compared to that in control cones (Fig. 5A, B; see also Fig. 5C). In addition, like the case in rods, cone sensitivity was also slightly reduced in Kv8.2 KO cones compared to controls (Fig. 5C, inset). Also, in resemblance to rods, mutant cone responses recovered 53% faster than those of control cones (Fig. 5D; Table 2). In summary the *in vivo* whole retina response to activation of rods versus cones is different but in isolation, the loss of Kv8.2 affects rod and cone responses in the same way.

**Figure 5:**
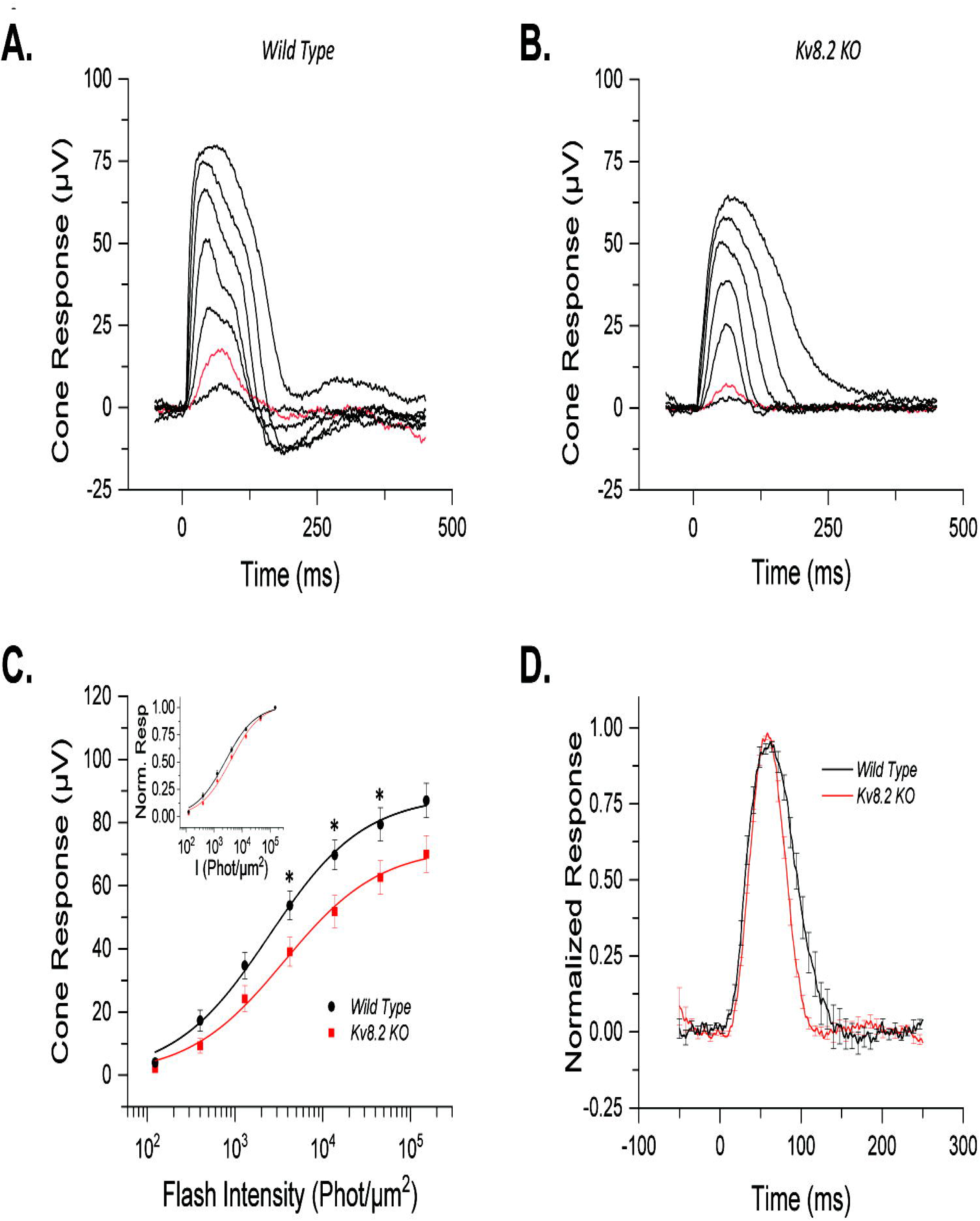
Ex-vivo responses of Kv8.2 KO cones. Representative responses to a family of flashes (123, 401, 1295, 4223, 13717, 45465 and 151808 photons) from ***A)*** wild-type control and ***B)*** Kv8.2 KO cones. The red traces in each case represent the dim flash response to 401 photons µm^-2^, highlighted for comparison. ***C)*** Averaged intensity response curves for Kv8.2 KO (red) and control (black) cones fit to a Naka-Rushton function; ***inset)*** normalized intensity response curves. ***D)*** Normalized dim flash response (red traces in ***A*** and ***B***) for Kv8.2 KO cones (red) and wild-type (black) cones.

**Table 2:**
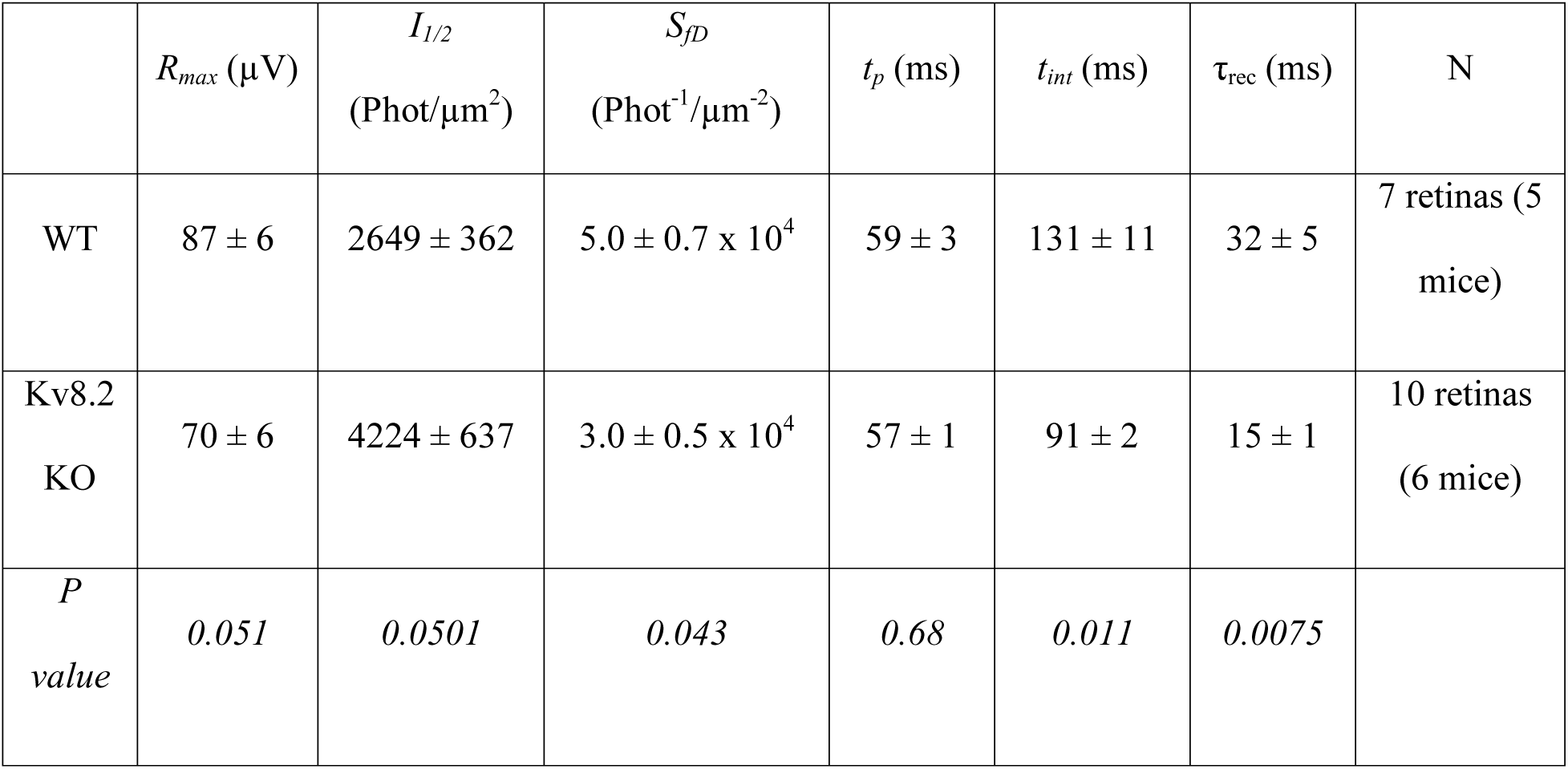
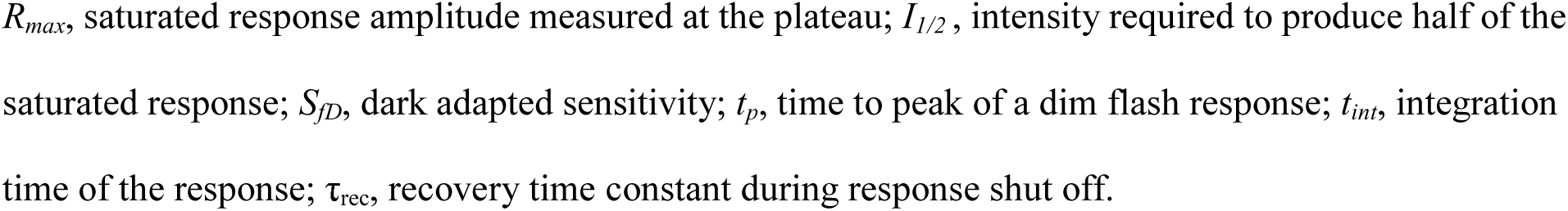
Cone response parameters from ex vivo recordings.

| | $R_{max}$ (μV) | $I_{1/2}$<br>(Phot/μm<sup>2</sup>) | $S_{\mathcal{D}}$<br>(Phot<sup>-1</sup>/μm<sup>-2</sup>) | $t_p$ (ms) | $t_{int}$ (ms) | $\tau_{rec}$ (ms) | N |
| --- | --- | --- | --- | --- | --- | --- | --- |
| WT | 87 ± 6 | 2649 ± 362 | 5.0 ± 0.7 x 10<sup>4</sup> | 59 ± 3 | 131 ± 11 | 32 ± 5 | 7 retinas (5 mice) |
| Kv8.2 KO | 70 ± 6 | 4224 ± 637 | 3.0 ± 0.5 x 10<sup>4</sup> | 57 ± 1 | 91 ± 2 | 15 ± 1 | 10 retinas (6 mice) |
| *P*<br>*value* | 0.051 | 0.0501 | 0.043 | 0.68 | 0.011 | 0.0075 |  |

$R_{max}$ , saturated response amplitude measured at the plateau; $I_{1/2}$ , intensity required to produce half of the saturated response; $S_{fd}$ , dark adapted sensitivity; $t_p$ , time to peak of a dim flash response; $t_{int}$ , integration time of the response; $\tau_{rec}$ , recovery time constant during response shut off.

### Retinal Degeneration

The Kv8.2 KO retina was thinner than WT at 3.5 months (Fig. 1D), so we next used OCT imaging to quantitate changes in gross retina anatomy from ages 2-10 months (Fig. 6). The outer nuclear layer (consisting of rod and cone nuclei) of Kv8.2 KO animals thinned progressively at a rate of 1.35 ± 0.11 µm/mon. At all ages, the ONL thickness in Kv8.2 KO animals was significantly reduced when compared to their wild type siblings (p < 0.001 at all ages, Sidak’s multiple comparison using mixed effect analysis) (Fig. 6B). That amount of ONL thinning corresponds to 10% loss at 2 months of age that progressed to 31% at 10 months of age in Kv8.2 KO mice (Fig. 6C). A common consequence of photoreceptor death is reactive Muller cell gliosis which can be measured as an increase in GFAP expression. We documented increased expression of GFAP in the Kv8.2 KO retina at 7 months of age by Western blot and immunohistochemistry (Fig. 6D, E).

**Figure 6:**
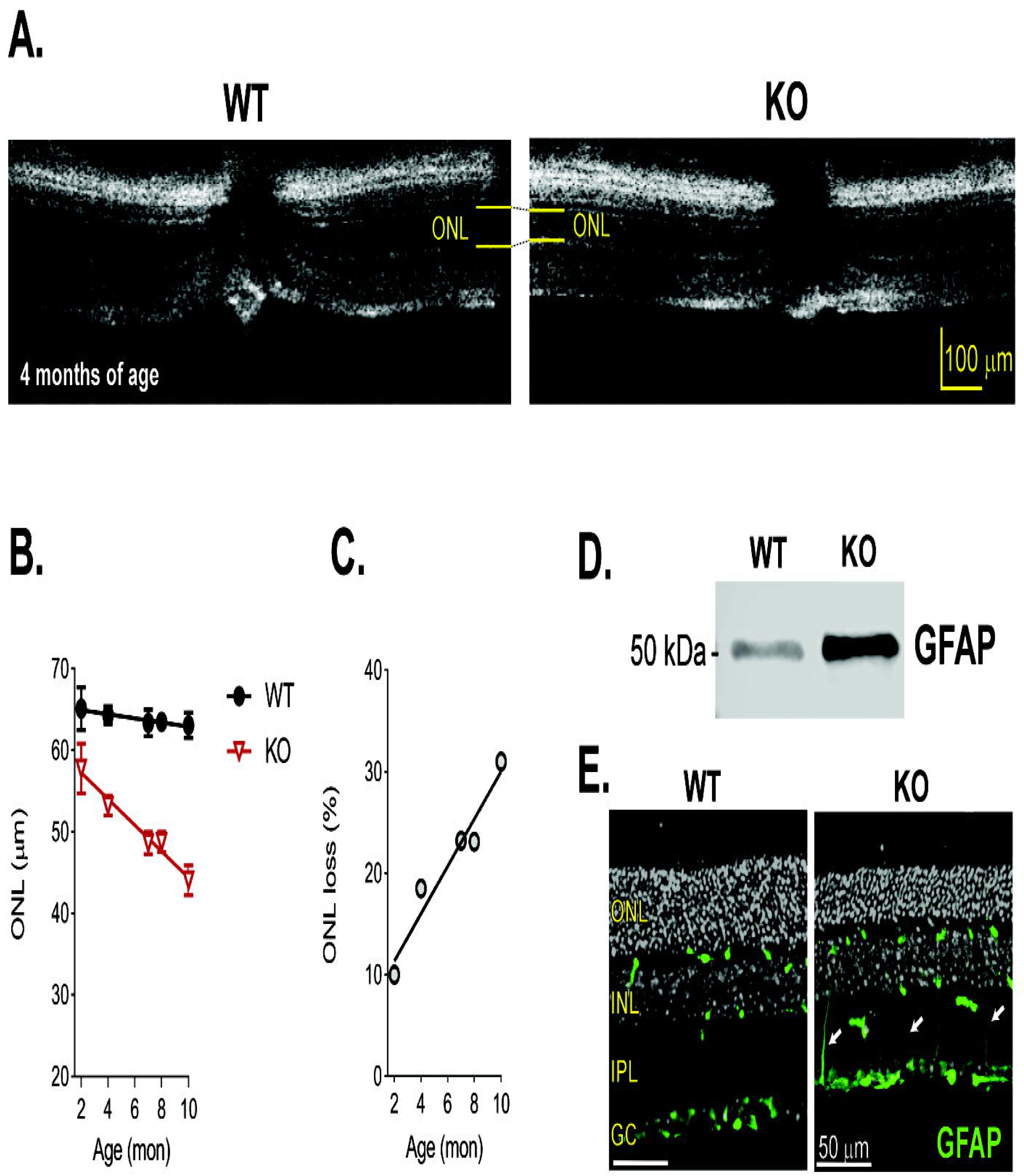
Kv8.2 KO retina undergo a mild retinal degeneration. ***A)*** OCT images of WT and Kv8.2 KO at 4 months of age with the thickness of the ONL layer indicated in yellow. ***B)*** ONL layer thickness in WT (black) or Kv8.2 KO (red) retina from ages 2-10 months. ***C)*** Percent thinning of the Kv8.2 KO ONL relative to that of WT. Upregulation of GFAP at 7 months is evident by both Western ***(D)*** and immunohistochemistry ***(E).*** Scale is 10 µm (*A*) or and 50 µm in (*E)*, abbreviations are ONL, outer nuclear layer; INL, inner nuclear layer; IPL, inner plexiform layer; and GC, ganglion cell layer.

The human macula is an area of high cone density, and the ERG results demonstrate that cone function is diminished in both humans and in our Kv8.2 KO mice. Therefore, even though the mouse retina does not have a macula, we expected an age-dependent loss of cones. Cone density was measured in retinal flatmounts immunolabeled with anti-cone arrestin. Initial analysis of cone density at younger ages did not reveal any changes (data not shown) so we aged animals to 12-months. There is a gradient of cone density in the mouse retina such that it is highest in the center (18). To account for this, we measured cone density at three different eccentricities from the optic nerve (Fig. 7A). The density of cones was indistinguishable between WT and Kv8.2 KO animals at all eccentricities (Fig. 7B; at center Δ mean= 506, 95% CI (-2813, 3825), one way ANOVA adj. p= 0.9737; at mid retina Δ mean= 198, 95% CI (-3121, 3517), one way ANOVA adj. p= 0.9983; at periphery Δ mean= -723, 95% CI (-4042, 2596), one way ANOVA adj. p= 0.9290).

**Figure 7:**
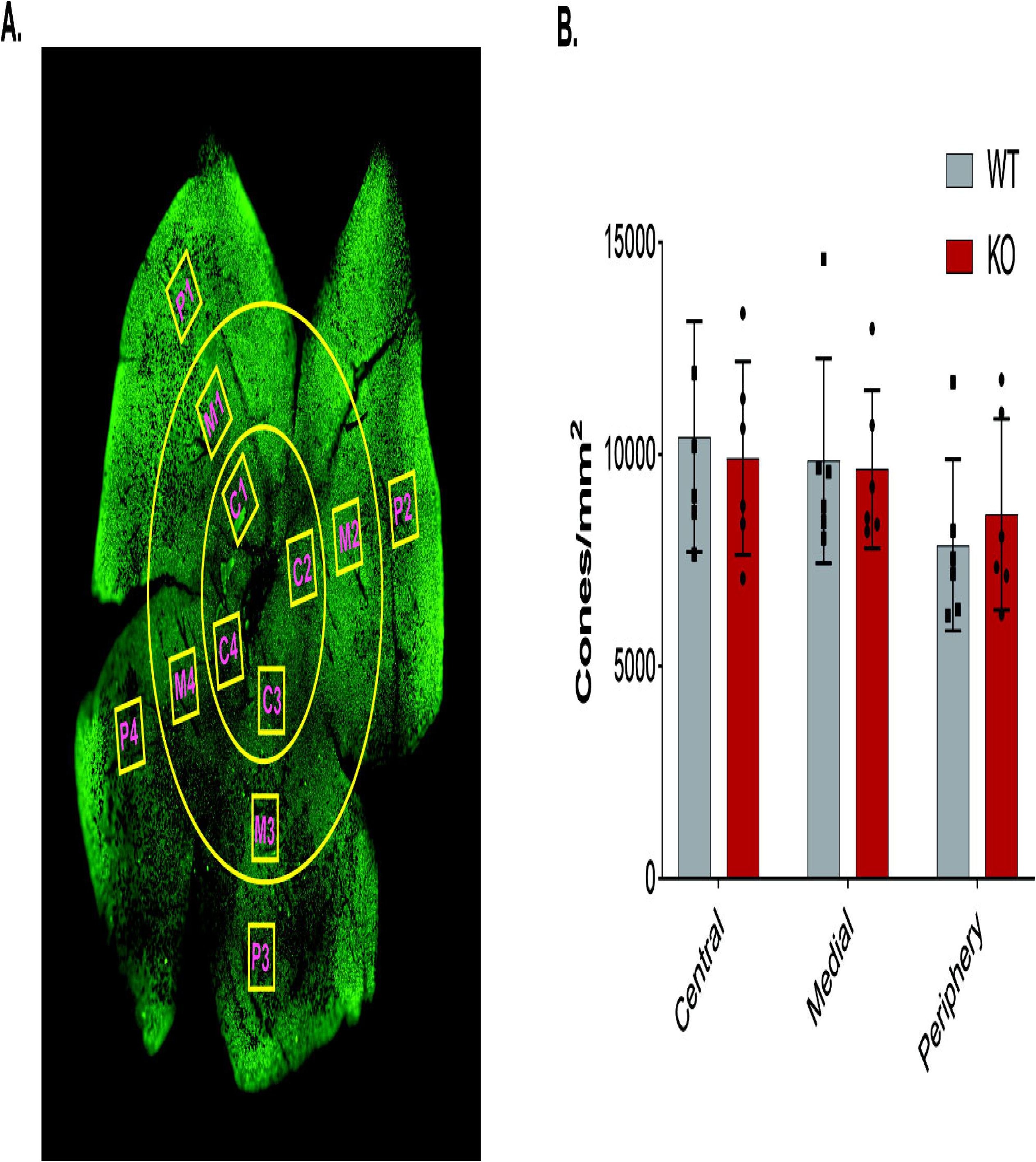
Cone density in Kv8.2 KO retina is comparable to its WT sibling at 12m of age. ***A)*** Representative image of a cone arrestin labeled retina showing the different ROI’s drawn at three different eccentricities from the optic nerve. ***B)*** Quantification of cone density as a function of retinal eccentricity.

We then assessed whether there were any ultrastructural differences between the WT and Kv8.2 KO retina at three-months of age. The organization of membranes in the outer and inner segment was normal (Fig. 8A, B). Dying photoreceptors in the form of autophagic-like or necrotic-like structures were found in the region of inner segments in the retina from Kv8.2 KO animals and sometimes membrane debris originating from both mitochondria and OS membranes were evident in these structures (Fig. 8C, D). In contrast to the Kv2.1 KO, we did not see any alteration in the number, appearance, or position of the mitochondria in the inner segments of mutant Kv8.2 KO photoreceptors (Fig. 8A-B, E-F).

**Figure 8:**
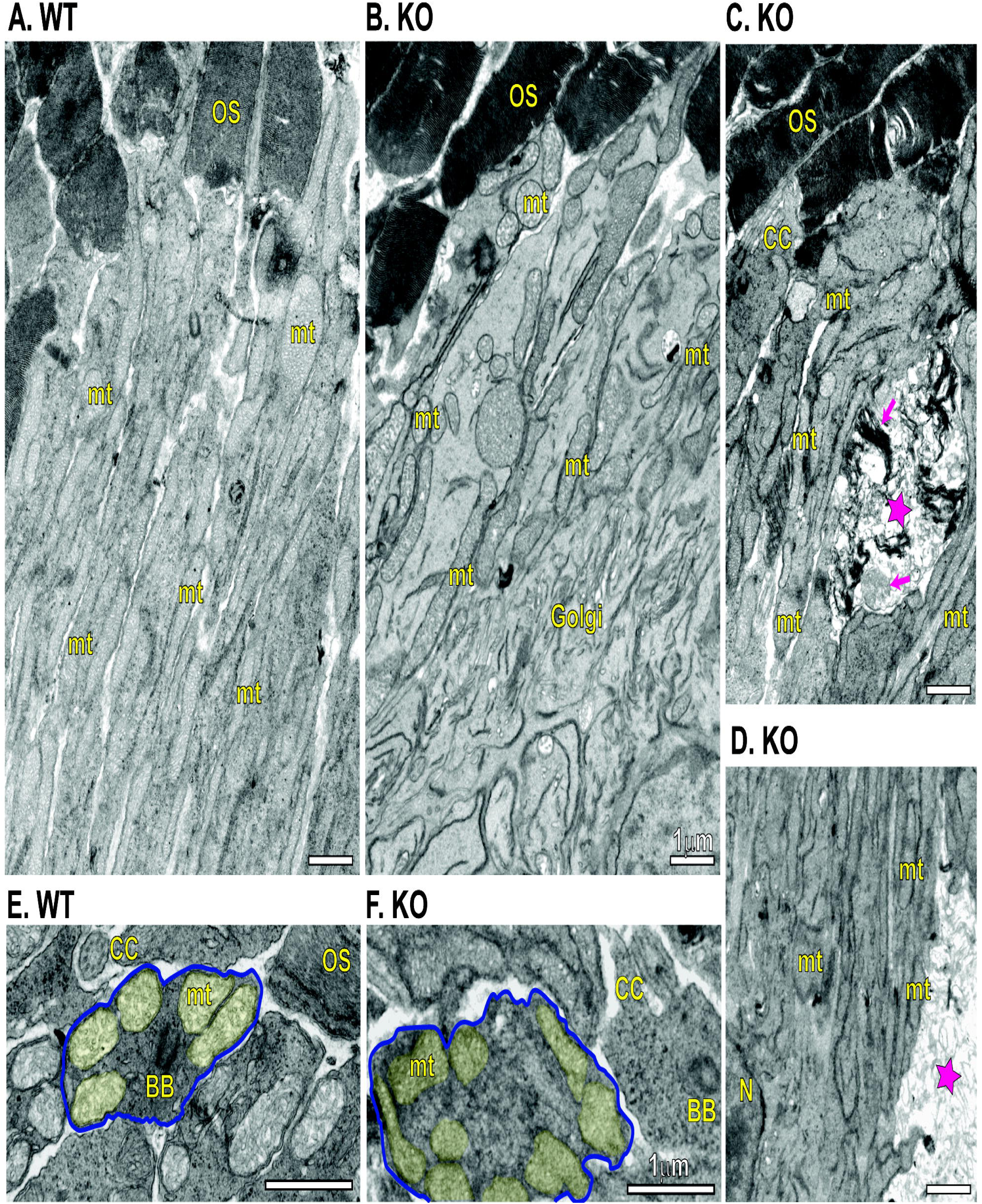
Ultrastructural comparison of WT and Kv8.2 KO photoreceptors. Inner segments from WT ***(A)*** and KO ***(B)***. ***C-D)*** Examples of inner segments from KO demonstrating presence of dead cell fragments, marked with magenta stars and magenta arrows indicating remnants of outer segment discs or mitochondria. Cross-section view of inner segments from WT ***(E)*** and KO ***(F)***, one cell in each is outlined in blue and mitochondria shaded in yellow. Scale bars are 1 µm and abbreviations are OS, outer segment; CC, connecting cilium; BB, basal body; mt, mitochondria; Golgi, Golgi Body; N, nucleus.

### Photophobia

Finally, we considered another common presentation of CDSRR which is photophobia. Photophobia can be measured in mice using a light-dark preference behavioral assay. Interestingly, photophobia has been reported in a mouse model of bradyopsia, another retinal disease characterized by altered timing of photoreceptor responses (19). We used the same light-dark preference assay to test Kv8.2 KO animals but found no evidence of light aversion (Supplemental Fig. 7).

### Kv8.2 KO in an all-cone retina

The ERG analysis demonstrated a reduction in cone function that cannot be attributed to cone loss. To better study cones, we crossed the Kv8.2 KO to a mouse strain with an all-cone retina C NRL KO; RPE65 R91W (20), that we refer to as “Conefull” to clearly distinguish this line from the original NRL KO. This line was chosen because it does not form the neural rosettes found in NRL KO that interfere with morphological analysis of the outer nuclear layer. his line provides a mouse anatomical equivalent to a cone-only fovea as is found at the center of the human macula. ERG analysis was carried out at between one and two months of age. Dark adapted animals were stimulated with test flashes increasing in intensity from -0.6 to -2.9 log (cd.s/m^2^). For descriptive statistics of this experiment, see Supplemental Table 9B and 9C. As expected, there was no response at the lower intensity flashes that activates rods. Beginning at the 1.9 log (cd.s/m^2^) flash, a cone-driven b wave was elicited. The cone-driven response was also present in the Conefull: Kv8.2 KO but reduced in amplitude with no change in the a wave (Fig. 9A-C). The histology of the Conefull: Kv8.2 WT and KO was similar at 2 months of age ( Δ mean= 15.5, 95% CI (-1.5, 32.5), two-way ANOVA adj. p= 0.0720). But by 4 months of age there was significant thinning of the outer nuclear layer in the KO compared to controls (Δ mean= 23.3, 95% CI (10.9, 35.7), two-way ANOVA adj. p= 0.0018) (Fig. 9D-E). This indicates that there is a progressive loss of cones in this unique retinal environment. These experiments demonstrate that the depressed cone signaling characteristic of CDSRR is intrinsic to altered cone signaling.

**Figure 9:**
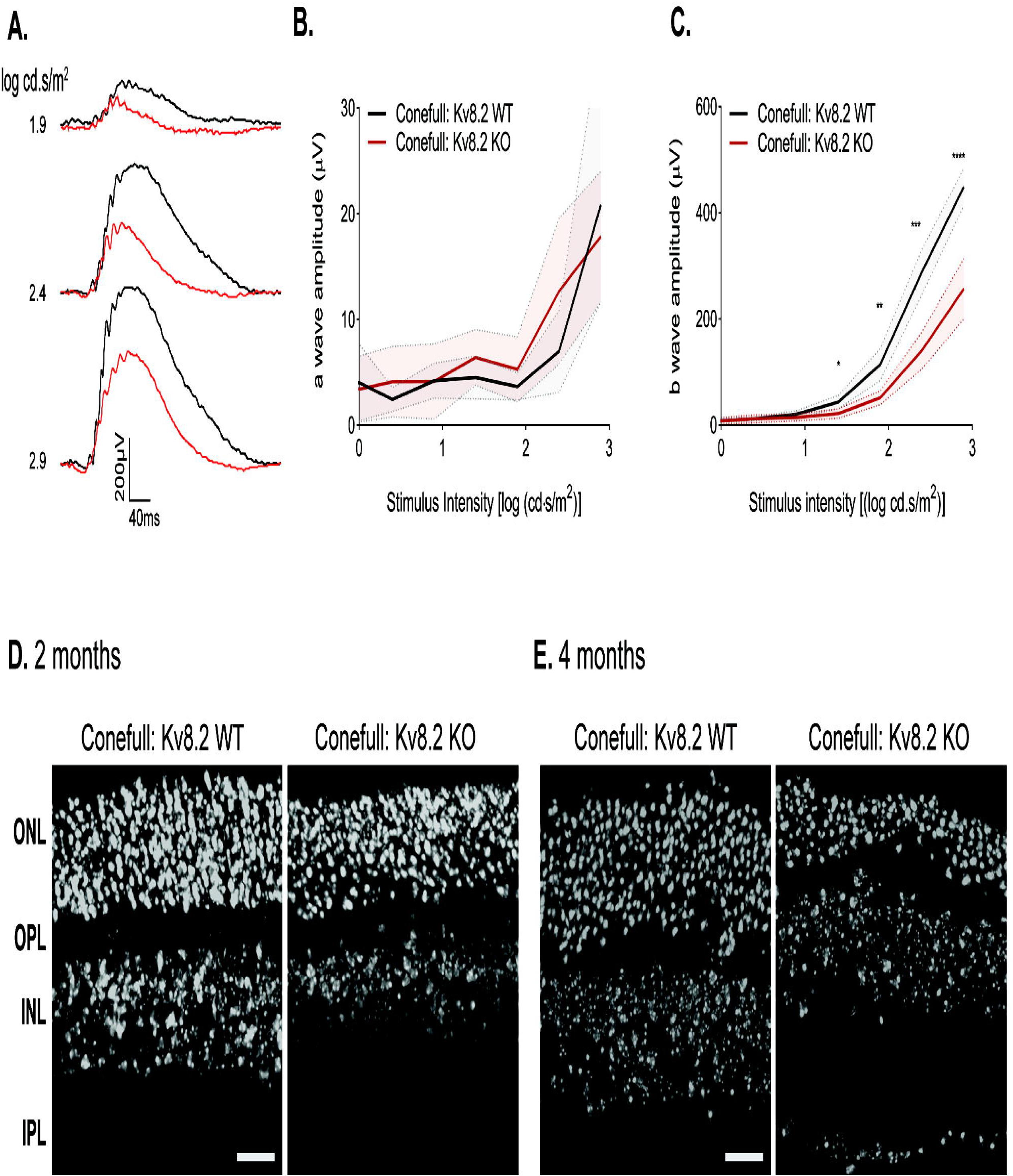
Kv8.2 KO in an all-cone retina. The all-cone retina mouse strain used here, “Conefull”, is the NRL KO combined with an RPE R91W knock-in. ***A)*** Representative ERG traces from the three brightest flashes in the photopic series using the ERG protocol in Fig. 8, Conefull: Kv8.2 WT (black) and Conefull: Kv8.2 KO (red). Scale is 200 µV by 40 ms. ***B)*** a wave and ***(C)*** b wave amplitude expressed as a function of flash intensity. ***C)*** Hoechst labeling of nuclear layers in retina sections from Conefull: Kv8.2 WT or Conefull: Kv8.2 KO at ***(D)*** 2 months or ***(E)*** 4 months of age. Scale is 20 µm in (*D,E)*, abbreviations are ONL, outer nuclear layer; OPL, outer plexiform layer; INL, inner nuclear layer; IPL, inner plexiform layer.

## DISCUSSION

The heteromeric voltage-gated K^+^ channel, Kv2.1/Kv8.2, participates in setting photoreceptor resting membrane potential. It carries most of the K^+^ efflux that balances the circulating dark current (9,10,13,21–23). Kv8.2 is lost or mutated in KCNV2 retinopathy, also called Cone Dystrophy with Supernormal Rod Responses (CDSRR) (2,3,8,14,24). Among inherited retinal degenerations, CDSRR stands out for its unique pattern of altered retinal signaling and early onset macular degeneration. The major question motivating this work was to determine how the phenotypes of the Kv8.2 KO mouse match what is reported for humans with CDSRR. Three major features of CDSRR are altered ERG responses, macular (cone) degeneration, and photophobia. The Kv8.2 KO mice only share the altered ERG phenotype. This work defines the limits of using mouse models to understand CDSRR and provides an in-depth analysis of Kv8.2-dependent rod and cone signaling.

### Comparison of the ERG defect in Kv8.2 KO mice and CDSRR patients

The signaling defects measured with ERG are similar between mouse and human. For simplicity we recommend comparing the data shown in Figure 2 with a recent review of CDSRR clinical presentations (4). In scotopic responses, the trough of the negative going a wave is square since onset of the positive going b wave is delayed. In humans the amplitude of the a wave is not appreciably altered whereas in the mouse, it is substantially reduced such that the supernormal b wave generated by brighter flashes of light can only be detected by normalizing to the a wave. The reduction in a wave amplitude in the mouse is seen in both Kv8.2 KO lines and Kv2.1 KO and is consistent with the reduced photoreceptor responses that we measured *ex vivo* (11–13). This agrees with the prediction that loss of Kv8.2 reduces the amount of circulating dark current so that light-evoked closure of CNG channels cannot generate as large of a response.

In CDSRR, a range of macular abnormalities are seen but only few changes outside of the cone-dense macula have been reported (25,26). The macula also has a high density of rods surrounding the cone-specific fovea, so at least in some CDSRR patients the progressive macular degeneration is likely to involve rods. The reduction in signaling in mice is close in magnitude, but not directly proportional, to the loss of rods. We propose a combination of curtailed circulating dark current with differing amounts of rod degeneration explains the variation in the ERG a wave of Kv8.2 KO mice and CDSRR patients.

### The role of Kv8.2 in rod function

We observed an unusual pattern in the timing of rod-driven photoresponses. At the dimmest flashes, rods generated a delayed but normal a wave amplitude. At brighter flashes the onset of the response was normal, but the amplitude was reduced. At all light intensities the rod-driven b wave was delayed. A slow or small response is logical given the expected reduction in circulating dark current, i.e. reduced [K^+^] in the extracellular subretinal space as described for the Kv2.1 KO (13). In support of this concept, our *ex vivo* ERG data, where any changes in the extracellular [K^+^] is washed out, did not show any delay in the onset of the response. Why the *in vivo* speed and magnitude of the response are differently affected as a function of light intensity will be an interesting area of future investigation.

The retina is most often exposed to complicated light patterns, so we extended our ERG analysis to include a sinusoidal, mesopic flicker and found a complex phenotype. The fundamental component of the response was attenuated at all frequencies in the Kv8.2 KO. Because of the complexity of this waveform and strikingly different response shapes formed by the WT and Kv8.2 KO at low frequencies, using the amplitude of the f_0_ may not reflect the true response amplitude. To circumvent this issue, we examined the amplitude of the b wave-like and c wave-like components at low frequencies. The b wave-like component was elevated in response to a 0.5 Hz flicker but attenuated at higher frequencies while the c wave-like component was reduced at all frequencies. The b wave-like response is similar to that observed using a repetitive mesopic flash flicker where the supernormal b wave is lost at frequencies higher than 0.5 Hz. The c wave-like component likely has the same origin as the c wave, which is a response of RPE and Muller glia to light triggered decreases of [K^+^] in the subretinal extracellular space (27–29). Thus the reduced c wave-like component is consistent with the attenuated c wave observed in Kv2.1 KO (13). In a separate study, the attenuation of the c wave in Kv8.2 KO, Kv2.1 KO, and Kv8.2/Kv2.1 double KOs was much larger so that component was essentially lost (12). The difference could reflect different ages of mice with corresponding different levels of gliosis. While further investigation into the origin of the waveforms elicited by the sinusoidal mesopic flicker will be necessary, we can conclude that loss of Kv8.2 impairs the response to repetitive stimulation in a frequency-dependent manner.

### The role of Kv8.2 in cone function

In CDSRR and Kv8.2 KO mice, cone-driven signaling is reduced. The simplest explanation for that could be the loss of cones as the macula degenerates. In support of this, Carvalho and colleagues reported a reduction in cone density in the Kv8.2 KO retina at six-months of age (12). However, this is one of the results from our independent lines of Kv8.2 KO that differ – we did not observe any cone loss in our Kv8.2 KO mice even at twelve-months of age. This difference likely originates in the smaller sampling of cones (∼5,500 vs our ∼10,000 cells/mm^2^). In the all-cone mouse retina, which we refer to as “Conefull” simply to clearly distinguish it from the parent NRL KO strain, we found that absence of Kv8.2 caused the photopic flash ERG responses to be reduced to the same magnitude as in the normal rod-dominant retina. Disappointingly, the Conefull retina did not respond to flicker ERG, possibly due to the rewiring of rod bipolar cells previously documented in the NRL KO (30). Nevertheless, we can conclude that the reduced cone driven signaling in the absence of Kv8.2 is intrinsic to altered cone physiology rather than being driven by coupling to rods.

To our surprise, we found that the outer nuclear layer of the Conefull: Kv8.2 KO thinned with age at a greatly accelerated rate compared to the Conefull controls. This observation will require further exploration and validation because while the Conefull (and parental NRL KO) line is a useful tool to study murine cones, the cones are not identical to cones in a normal retina. They are derived from S-cones, have aberrant synaptic wiring, and the Conefull strain does not form neural rosettes but still undergoes an early onset retinal degeneration. All the same, if cones with altered signaling are more susceptible to loss when they are not protected by surrounding rods, this could explain the selective macular degeneration in CDSRR.

### The potential role of Kv8.2 in regulating photoreceptor Kv2.1-containing channels

We recognize that loss of Kv8.2 does not only affect *I*_Kx_, but could alter the resting membrane potential and/or ion homeostasis, neither of which can be directly measured by ERG. In our opinion, the initial focus should be on how loss of Kv8.2 affects Kv2.1 function as that will set a baseline to ask more specific questions about how HCN1, or other ion channels in the retina are regulated by Kv2.1/Kv8.2 activity. Without modulation by Kv8.2, its binding partner Kv2.1 would instead form homomeric Kv2.1 channels. Kv2.1 homomeric channels are found throughout the body, independent of Kv8.2 or other KvS subunits, so it is not surprising that loss of Kv8.2 only resulted in a small increase in expression without altering the subcellular localization of Kv2.1. Carvalho and colleagues also reported that Kv2.1 localization is unaffected by loss of Kv8.2 although they report a decrease in Kv2.1 expression levels (11). Measuring expression levels by immunohistochemistry rather than Western blotting as we did is more prone to artifacts, but another important difference in our two independent Kv8.2 KO lines is the presence of *Kcnv2* transcripts in our line. It is not known if the transcripts for Kv8.2 and Kv2.1 have any interaction but if there is one, it could serve to provide regulation of mRNA stability or translation thus ultimately influencing the protein levels of Kv2.1. The mechanisms of coordinated regulation between Kv2.1 and Kv8.2 requires further investigation as there are several interesting phenomena noted in the literature. Including the circadian control over the expression of these channel subunits (31) and observations that Kv8.2 upregulation in the hippocampus influences susceptibility to epilepsy (32).

There is electrophysiological evidence for the presence of Kv2.1 homomeric channels even in healthy rods and such channels activate at more positive potentials than the normal resting membrane potential of photoreceptors (9). If Kv2.1 homomeric channels were inactive in the Kv8.2 KO retina, we would expect to see the same phenotypes in the Kv8.2 and Kv2.1 KO mice. Alternatively, if Kv2.1 homomeric channels were active in the Kv8.2 KO retina we would expect to see differences in the electrical properties between the two mutant mice lines. The two models are similar, but not identical. Most notably, the inconsistency in the reports of a supernormal b wave response in Kv2.1 KO mice needs to be further investigated (11–13). Both lines of mice exhibit mild degeneration, and the mechanism in the Kv2.1 KO retina involves increased calcium influx via CNG channels and mitochondrial dysgenesis (13). Our data on Kv8.2 KO mice do not directly address that issue. However, the normal appearance of mitochondria in Kv8.2 KO EM images indicates that the mechanism of cell death is different from what occurs due to loss of Kv2.1. Future direct recordings of photoreceptors from WT, Kv8.2 KO, and Kv2.1 KO performed side by side could unravel the mechanism of these differences.

In conclusion, we describe the defects in an independent Kv8.2 KO mouse which has helped us to understand that CDSRR is a result of complicated interactions between altered K^+^ homeostasis differentially affecting the magnitude and timing of photoresponses, and mild degeneration affecting rods more than cones. Kv8.2 KO mice are a valuable resource for continued studies of photoreceptor physiology.

## MATERIALS AND METHODS

### Animals

Kv8.2 KO mice were created at the Genome Editing Facility at the University of Iowa. C57Bl6/J zygotes were electroporated with Cas9, tracrRNA, and 2 crRNA (5’ CCCCAGAGACTGAGGATGTGCCC and 5’ CCTCGGTCCATTGTGGTGCGATG). Five of the 25 resulting animals with homozygous deletions were sequenced at the *Kcnv2* locus and two with homozygous 101 bp deletions were selected as founders. Both founder lines (3651.4 and 3654.1) were backcrossed to C57BL/6J (RRID:IMSR JAX:000664) for 2 generations. No significant differences were found in the data obtained from the different founder lines but for convenience the bulk of the experiments were conducted with animals from the 3654.1 founder. The colony was maintained by crossing heterozygote siblings and resulting KO animals were analyzed in parallel with the WT siblings for controls. NRL KO; RPE65 R91W; (Conefull) mice were a generous gift from Christian Grimm (20), Animals were genotyped by PCR (KCNV2 primers, 5’ ATAGACAGGCAGGAGAGATAGG and 5’ TCACAGCATTCCAGCAGATAG, generate a 410 or 309 bp product for the WT or KO allele respectively) or via the services of Transnetyx (Cordova, TN). Mice of both sexes, between the ages of 1- 14 months were used. Mice were housed in a central vivarium, maintained on a standard 12/12-hour light/dark cycle, with food and water provided *ad libitum* in accordance with the Guide for the Care and Use of Laboratory Animals of the National Institutes of Health. All procedures adhered to the ARVO Statement for the Use of Animals in Ophthalmic and Vision Research and were approved by the University of Iowa IACUC committee.

### RNA preparation and (Reverse Transcriptase-digital droplet PCR) RT-ddPCR

Performed and analysed as previously described (33). The probe sets for mouse KCNV2 (Mm.PT.58.6815878) was purchased from IDT (Coralville, IA).

### Western Blotting

Retinas were collected by dissection from 3-month old animals. Sets of two retinas were lysed in 250 µl lysis buffer (50 mM Tris pH 7.5, 150 mM NaCl, 5 mM EDTA, 1% Triton-X-100) supplemented with 1x protease inhibitor (Complete mini EDTA-free, Roche, Switzerland and 1x phosphatase inhibitor (PhosStop, Roche, Switzerland) on ice. Samples were clarified by centrifugation at 16,873 x g at 4 °C for 15 minutes. Protein concentration was measured using the BCA assay (Thermo Scientific, Massachusetts, USA). Reducing sample buffer was added, samples heat denatured and 20 µg of total protein was loaded per lane on 4-20% Tris-HCl polyacrylamide gels (BioRad, California, USA). After transfer to PVDF, the following primary antibodies were used for blotting: Kv8.2 N448/88 at 1:1000 (Antibodies Incorporated Cat# 75-435, RRID:AB_2651161)), Kv2.1 K89/34 at 1:1500 (Antibodies Incorporated Cat# 75-014, RRID:AB_10673392), rabbit anti-HCN1 at 1:3000 (34), and Na/K-ATPase M7-PB-E9 at 1:1000 (Santa Cruz Biotechnology Cat# sc-58628, RRID:AB_781525)). Goat anti-mouse or rabbit antibodies were conjugated to IRdye 680 or 800 (LI-COR Biosciences Cat# 925-32210, RRID:AB 2687825, Cat# 925-68071, RRID:AB_2721181). Blots were imaged using a LI-COR Odyssey FC and analyzed with Image Studio (v5) software. Signal intensities were normalized to that of total protein which was imaged using Revert total protein stain (Li-Cor) prior to blotting.

### Immunohistochemistry

Immunostaining was carried out as previously described (34). Briefly, posterior eyecups were collected by dissection, fixed in 4% paraformaldehyde (PFA) at room temperature for 15- 60 min, cryoprotected in 30% sucrose, and then frozen in Tissue-Tek (Electron Microscopy Sciences, Hatfield, PA). Radial sections were cut and collected on electrostatically charged glass slides, and either labeled immediately or stored at -80 °C until use. Blocking buffer consisted of 10% normal goat serum and 0.5% Triton X-100 in PBS. Antibodies diluted in blocking buffer were incubated on retinal sections for 1-3 h at room temperature or overnight at 4° C. Primary antibodies used were: Kv8.2 N448/88 at 1:1000 (NeuroMab, AB_2651161), Kv8.2 N448/50.1 at 1:50 (gift from James Trimmer), Kv2.1 K89/34 at 1:1500 (NeuroMab, AB_10673392), and cone arrestin at 1:500 (Millipore Cat# AB15282, RRID:AB_1163387). Secondary antibodies were conjugated to Alexa 488, 568, or 647 (Thermo Fisher Scientific Cat# A-11001, RRID:AB_2534069, Cat# A-11004, RRID:AB_2534072, or Cat# A-21235, RRID:AB_2535804) and nuclei were stained with Hoechst 33342 (Life Technologies, Carlsbad, CA) and mounted with Fluoromount-G (Electron Microscopy Sciences, Hatfield, PA). Images were collected on a Zeiss LSM710 confocal (Central Microscopy Research Facility, University of Iowa) or Leica DM6B, THUNDER Imager. For image analysis, maximum through z-stack projections were used with manipulation limited to rotation, cropping, and adjusting the brightness and contrast levels using Image J, Zen Light 2009 (Carl Zeiss, Oberkochen, Germany) or Photoshop CC (Adobe, San Jose, CA). A minimum of two images per mouse for at least two mice per genotype per experiment were analyzed.

### Electroretinography (ERG)

All ERG recordings were obtained on the Espion V6 Diagnosys Celeris system (Diagnosys LLC, Lowell, MA). Mice were dark-adapted overnight, and all steps performed on the day of ERG recording were done in dim red light. Mice were anesthetized with a mixture of ketamine (87.5 mg/kg) and xylazine (2.5 mg/kg). Tropicamide (1%) was used to dilate the pupils and Genteal or Systane gel (0.3% Hypromellose) (Alcon, Geneva, Switzerland) was used to keep the eyes hydrated. Body temperature of the mice was maintained by keeping them on a heating pad. Five different protocols were used. The scotopic and photopic series protocols were performed using a reference electrode placed subcutaneously along the nasal ridge and grounding electrode placed subcutaneously into the haunch. The International Society for Clinical Electrophysiology of Vision (ISCEV) and two flicker protocols were performed without a reference or ground by alternating eye stimulation and using the non-stimulated contralateral eye as a reference. Mice ranged in age from 1-5 months, with 9 WT littermate controls and 8-9 Kv8.2 KO animals. Conefull animals were tested at 1 month of age using 6 Conefull: KCNV2 WT littermate controls with 13 Conefull: Kv8.2 KO mice.

The ISCEV protocol (15) as applied to mice, consisted of 5 steps, 2 recorded from the dark-adapted mice and the latter three after 10 minutes of light-adaptation (9 cd.s/m^2^): 1) scotopic dim flash (-2 log cd.s/m^2^), 2) scotopic bright flash (0.5 log cd.s/m^2^), 3) standard combined response to a photopic flash (0.5 log cd.s/m^2^), 4) 5 Hz flicker, and 5) 30 Hz flicker. 5-20 responses were recorded at each step. The scotopic intensity series consisted of 8 steps using a white flash ranging in intensity from -2.5 to 2 log cd.s/m^2^. For intensities from -2.5- -1 log cd.s/m^2^, 10 responses were recorded at intervals of 5 s, for 0 and 0.5 log cd.s/m^2^ 15 responses were recorded at intervals of 10 s, for 1 and 1.5 log cd.s/m^2^ flash, 5 responses were recorded at intervals of 10 s, after 30 s a single recording was collected at the highest intensity (2 log cd.s/m^2^). For the photopic intensity series, the animals were first light adapted for 10 minutes and background illumination (1.5 log cd.s/m^2^) was maintained as the responses to a series of increasingly bright test flashes (-0.6, 0.0, 0.4, 0.9, 1.4, 1.9, 2.4, 2.9 log cd.s/m^2^) were recorded. Recordings were collected from both left and right eyes, individual responses at each step in each respective protocol was averaged. The a-wave amplitude was measured as the amplitude from the baseline to the first negative trough. The b-wave amplitude was measured from the peak of the a-wave to the third positive peak. Implicit times were measured from the onset of stimulus to the peak amplitudes of a- and b- waves respectively.

The repetitive flash flicker series consisted of 12 steps using a 0.5 log cd.s/m^2^ flickering white light with increasing frequencies (0.5, 1, 2, 3, 5, 7, 10, 12, 15, 18, 20, and 30 Hz). Responses were limited to 300ms and were averaged from 3 recordings for 0.5-3 Hz, 5 recordings for 7 Hz, 20 recordings for 10-15 Hz, and 50 recordings for 18-30 Hz. There was no delay between recordings and the first recording was always rejected. Amplitudes were measured from the trough to the peak. The sinusoidal flicker series consisted of an initial 10 minute adaptation step to a 0 log cd/m^2^ white background light followed by sinusoidal modulation with 100% contrast at increasing frequencies (0.5, 0.75, 1, 2, 3, 6, 9, 12, 15, and 25 Hz). Responses were averaged from 20 recordings for frequencies from 0.5-12 Hz and 50 recordings for higher frequencies. Responses were limited to 4 s for frequencies from 0.5-1.5 Hz, 2 s for frequencies from 2-3 Hz, 1 s for frequency of 6 Hz, and 300 ms for frequencies from 9-25 Hz. There was no delay between recordings and the first recording was always rejected. Responses were quantified by multiple means. The magnitude of the fundamental was assessed by applying the fast Fourier transform using MATLAB (MathWorks R2021a). The amplitude of the b wave-like and c wave-like peaks were measured from the trough of the response, the absolute minima, to the peak of each component.

### Photophobia

The light aversion assay was performed as described in (19). All animals were tested between 8 and 10 weeks of age. Light adapted animals were acclimated to the testing room for one hour prior to analysis. Testing chambers were divided into equal sized light and dark zones with a small opening between to allow animals to freely move between the two chambers. Animal movement was monitored by infrared rays. White light was provided by an LED array above the testing chamber, giving an illumination of 25,000 lux. Naïve animals were placed in the light zone and their movement was recorded for 20 minutes. Sample size was 14 WT and 15 Kv8.2 KO animals.

### Ex vivo ERG

The animals were dark-adapted overnight prior to the experiment and were euthanized by incubating in a CO_2_ chamber in darkness. The eyes were enucleated in dim red light and then transferred to a Petri dish containing oxygenated Ames medium (Sigma-Aldrich, St. Louis MO) at room temperature. Dissection was performed under infrared illumination using a microscope. The eyeballs were hemisected along the ora serrata to separate the cornea and lens from the posterior eyecup. The retina was then gently detached from sclera and RPE using forceps and stored in oxygenated Ames medium in a dark chamber until recording. Recordings were done using previously described methods (35). Briefly, the retina was mounted in a closed chamber with the photoreceptors facing up and placed under a microscope (Olympus IX51). The recording chamber was supplied with heated Ames medium at a flow rate of 3-5 ml/min. To isolate the photoreceptor component from the retinal response, 50 μM DL-AP_4_ (Tocris Bioscience, Bristol United Kingdom) and 100 μM BaCl_2_ (Sigma-Aldrich, St. Louis MO) were included in the Ames medium. The chamber temperature was closely maintained at 35-36 °C. The retinas were adapted to the recording conditions for at least 15 minutes before experiments. Ex-vivo transretinal recordings were made by presenting a family of 530 nm light flashes. The flashes were generated by computer-controlled LEDs (Thor Labs, Newton, NJ) and guided through the microscope optics onto the retina. The flash responses were amplified using a differential amplifier (Warner Instruments, Hamden, CT), low-pass filtered at 300 Hz (Krohn Hite Corp., Brockton, MA), digitized using Digidata 1440 (Molecular Devices, San Jose, CA), and sampled at 10 kHz. Data was acquired and recorded on a computer using pClamp 10 software. To separate the cone response from the rod response, first a probe flash (10,000 photons µm^-2^) was presented to saturate the rod response. Then after 350 ms, a series of flashes was presented to record cone response as described earlier (35). Data was analyzed using pClamp 10, Microsoft Excel and Origin 9.8.5 (64 bit, SR2, OriginLab) and presented as mean ± SEM. Flash family response curves were fit to a Naka-Rushton function using the equation:

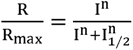

Where, R_max_ is the maximum response amplitude, I is the flash intensity, n is the Hill coefficient and I_1/2_ is the intensity to produce half-saturating response.

### Optical Coherence Tomography (OCT)

Mice were anesthetized with a mixture of ketamine (87.5 mg/kg) and xylazine (2.5 mg/kg). Tropicamide (1%) was used to dilate the pupils and Genteal (0.3% Hypromellose) (Alcon, Geneva, Switzerland) was used to keep the eyes hydrated. Images were collected with a Bioptigen spectral-domain imaging system (Bioptigen, Inc.) equipped with a mouse retina objective, reference arm position set at 1264. Scan parameters were as follows: rectangular (1.4 mm^2^) volume scans, 1000 A-scans/B-scan, 33 B-scans/volume, 3 frames/B-scan, and 1 volume. To quantify degeneration rates, the distance between the OLM and OPL bands was measured using Photoshop CC (Adobe) with calibration to the vertical scale bar in 4-8 images adjacent to the optic nerve for each animal; 4-6 animals were used for each genotype at each age.

### Cone density measurements

The edges of whole retinas were nicked to allow the retina to lay flat and placed on a nitrocellulose membrane (0.8 µm, Millipore, MA, USA) with the photoreceptor side up. Flatmounts were fixed in 4% PFA at room temperature for 30 minutes, rinsed in PBS and blocked in 5% goat serum with 0.1% TritonX-100 in PBS for 1 hour. Flatmounts were incubated in anti-cone arrestin 1:500 (Millipore Cat# AB15282, RRID: AB_1163387) overnight, rinsed and incubated with goat-anti rabbit: Alexa488 diluted 1:200 (Thermo Fisher Scientific Cat# A-11001, RRID:AB_2534069) for 1 hour. Images were acquired on a Leica DM6B, THUNDER Imager. A tiled image of the entire flatmount was first collected at 10x magnification. Two concentric circles were drawn centered on the optic nerve to mark the central, medial, and peripheral eccentricities of the retina. Within each eccentricity and each leaf of the retina, a region of interest (ROI) covering the same area was drawn (3 ROI’s per leaf). 40x images of each ROI were analyzed in Image J using the analyze particle function after thresholding the images. Cone density within the same eccentricity and animal were averaged. Comparisons were made using 2- way ANOVA and Sidak’s multiple comparisons test. Six animals of each genotype at 12-months of age were used.

### Transmission electron microscopy

Posterior eye-cups, collected from 3-month old WT and Kv8.2 KO mice, were fixed with a mixture 2% PFA and 2% glutaraldehyde in a buffer containing either 0.1 M sodium cacodylate, pH 7.4 (Univ. of Iowa) or 50 mM MOPS, pH 7.4, 0.05% calcium chloride (UC- Davis) then stained, embedded in Epon and sectioned as previously described (13,36).

### Experimental Design and Statistical Analysis

Statistical differences were determined using GraphPad Prism software (v8) Statistical significance was defined using an alpha of 0.05. T-tests, repeated measures ANOVA or mixed model effects. The specific test used is indicated in the results section. Sidak’s post-hoc test was used for multiple comparisons. Unless indicated otherwise, mean is reported with the SEM and variability is shown in all graphs by plotting the SD. P-values are indicated with asterisks where, *p < 0.05, **p < 0.01, ***p < 0.001, and ****p < 0.0001. Complete descriptive statistics are provided in supplementary tables.

## Supporting information

Supplemental Statistical Tables

## ACKNOWLEDGEMENTS

We thank William Paradee, Director of the University of Iowa Genome Editing Facility for his help designing the strategy to generate the Kv8.2 KO mice. We also thank James S. Trimmer from UC-Davis who generously provided previously untested mouse monoclonal anti-Kv8.2 supernatants. And we appreciate Andrew Russo from the Department of Molecular Physiology and Biophysics at the University of Iowa for use of his equipment to perform the light aversion assay. We received excellent advice from Ying Guo and Yumiko Umino at SUNY Upstate Medical University regarding the analysis of flicker ERG. We also thank Marie Burns at UC-Davis for helpful discussions and reading of the manuscript.

This work was supported by the National Eye Institute (R01 EY020542 to SAB, R01s EY027387 and EY030912 to VJK, R01 EY026216 to ES) and Foundation Fighting Blindness (BR-CMM-0619-0763- UIA to SAB). We also acknowledge unrestricted grants from Research to Prevent Blindness to the Department of Ophthalmology and Visual Sciences at SUNY Upstate Medical University, NY and the Department of Ophthalmology at University of California, Irvine, CA. We used the Zeiss LSM710 confocal in the University of Iowa Central Microscopy Research Facilities that was purchased with funding from NIH, SIG S10 RR025439. We also used the services of the Genomics Division of the Iowa Institute of Human Genetics which is supported, in part, by the University of Iowa, Carver College of Medicine.

## CONFLICT OF INTEREST STATEMENT

The authors declare no competing financial interests.

**Supplemental Figure 1 (related to Fig. 1):** Retinal transcripts for *Kcnv2* were reduced but not absent in the Kv8.2 KO mice. The amount of Kcnv2 in cDNA (0.25-20 ng) prepared from WT (black) or Kv8.2 KO (red) retina was measured by RT-ddPCR and fit with a simple linear regression, the slopes were significantly different (15.0 ± 0.4 for WT and 7.2 ± 0.1 for KO, p < 0.0001).

**Supplemental Figure 2 (related to Fig. 1):** Comparison of Kv8.2 antibodies for retinal immunohistochemistry. ***A***) Kv8.2 antibody clone N448/88 staining (green) shows presence of Kv8.2 protein in the IS of both WT and Kv8.2 KO retinas (non-specific staining). ***B)*** Kv8.2 KO antibody clone N488/50 shows a complete absence of Kv8.2 signal in the IS compared to the WT. Scale bar is 10 μm.

**Supplemental Figure 3 (related to Fig. 2):** Quantitation of ERG a and b wave amplitudes obtained from the ISCEV protocol shown in Fig 2. In each graph, WT mice are denoted by black bars and Kv8.2 KO by red bars ***A*)** Response of dark-adapted mice to a dim flash (-2 log cd.s/m^2^). ***B*)** Response of dark-adapted mice to a bright flash (0.5 log cd.s/m^2^). ***C*)** Standard combined response of light-adapted mice to a bright flash (0.5 log cd.s/m^2^). In all, the left panel is the a wave amplitude, center panel is the b wave amplitude and the right most panel is the ratio replotted from Fig 2 for reference. Plotted is the mean with the standard deviation.

**Supplemental Figure 4 (related to Fig. 2):** *Scotopic ERG response of Kv8.2 KO animals. **A)*** Representative ERG traces of dark-adapted WT (black) and Kv8.2 KO (red) mice to a single flash stimulus of increasing light intensity (log -2.5-2.0 cd.s/m^2^). ***B)*** a wave and ***(D)*** b wave amplitude expressed as a function of flash intensity. ***C)*** *a wave and **(E)** b wave* implicit time (time to peak) expressed as a function of flash intensity. ***F)*** Ratio of b wave to a wave amplitude. ***G)*** Implicit time of b wave with the implicit time of a wave subtracted. In all graphs, the scale is 200 µV by 40 ms, line is the mean, and the shaded area is the SD.

**Supplemental Figure 5 (related to Fig. 2):** *Photopic ERG responses of Kv8.2 KO animals. **A)*** Representative ERG traces of dark-adapted WT (black) and Kv8.2 KO (red) mice to a single flash stimulus of increasing light intensity (log -0.6-2.9 cd.s/m^2^) under rod-saturating background light. ***B)*** a wave and ***(D)*** b wave amplitude expressed as a function of flash intensity. ***C)*** *a wave and **(E)** b wave* implicit time (time to peak) expressed as a function of flash intensity. ***F)*** Ratio of b wave to a wave amplitude. ***G)*** Implicit time of b wave with the implicit time of a wave subtracted. In all graphs, the scale is 100 µV by 40 ms, line is the mean, and the shaded area is the SD.

**Supplemental Figure 6 (related to Fig. 2):** *Flash flicker ERG reveals frequency dependent attenuation of response amplitude in Kv8.2 KO*. ***A)*** Representative ERG traces of WT (black) and Kv8.2 KO (red) to a repetitive 0.5 log (cd.s/m^2^) flash of increasing frequency from 0.5-30Hz against a dark background. ***B)*** Response amplitude plotted as a function of stimulus frequency.

**Supplemental Figure 7:** *Light aversion assay*. ***A)*** The amount of time WT (black) and Kv8.2 KO (red) mice spent in the light exposed chamber binned into four 5-minute intervals to encompass the 20-minute testing time.

**Supplementary Tables = 18** Statistical Tables (SEPARATE FILE)

## ABBREVIATIONS

CDSRR: Cone Dystrophy with Supernormal Rod Responses
KCNV2: Potassium voltage-gated channel modifier subfamily V member 2
KCNB1: Potassium voltage-gated channel subfamily B member 1
Kv8.2: Voltage-gated potassium channel subunit 8.2
Kv2.1: Voltage-gated potassium channel subunit 2.1
ISEV: International Society for Clinical Electrophysiology of Vision
ERG: electroretinography
OCT: optical coherence tomography
HCN1: Hyperpolarization and cyclic nucleotide- gated channel, family member 1
NKA: Sodium-potassium ATPase
RPE: Retinal pigment epithelium
OS: Outer segment
IS: Inner segment
OLM: Outer limiting membrane
ONL: Outer nuclear layer
OPL: Outer plexiform layer
INL: Inner nuclear layer
IPL: Inner plexiform layer
GC: Ganglion cell layer
CC: Connecting Cilium
BB: Basal body
N: Nucleus

## REFERENCES

1. Bocksteins, E. (2016) Kv5, Kv6, Kv8, and Kv9 subunits: No simple silent bystanders. J Gen Physiol, 147, 105–125.

2. Gouras, P., Eggers, H.M. and MacKay, C.J. (1983) Cone dystrophy, nyctalopia, and supernormal rod responses. A new retinal degeneration. Arch Ophthalmol, 101, 718–724.

3. Wu, H., Cowing, J.A., Michaelides, M., Wilkie, S.E., Jeffery, G., Jenkins, S.A., Mester, V., Bird, A.C., Robson, A.G., Holder, G.E. et al. (2006) Mutations in the gene KCNV2 encoding a voltage-gated potassium channel subunit cause “cone dystrophy with supernormal rod electroretinogram” in humans. Am J Hum Genet, 79, 574–579.

4. Guimaraes, T.A.C., Georgiou, M., Robson, A.G. and Michaelides, M. (2020) KCNV2 retinopathy: clinical features, molecular genetics and directions for future therapy. Ophthalmic Genet, 41, 208–215.

5. Michaelides, M., Holder, G.E., Webster, A.R., Hunt, D.M., Bird, A.C., Fitzke, F.W., Mollon, J.D. and Moore, A.T. (2005) A detailed phenotypic study of “cone dystrophy with supernormal rod ERG”. Br J Ophthalmol, 89, 332–339.

6. Robson, A.G., Webster, A.R., Michaelides, M., Downes, S.M., Cowing, J.A., Hunt, D.M., Moore, A.T. and Holder, G.E. (2010) “Cone dystrophy with supernormal rod electroretinogram”: a comprehensive genotype/phenotype study including fundus autofluorescence and extensive electrophysiology. Retina, 30, 51–62.

7. Vincent, A., Wright, T., Garcia-Sanchez, Y., Kisilak, M., Campbell, M., Westall, C. and Heon, E. (2013) Phenotypic characteristics including in vivo cone photoreceptor mosaic in KCNV2-related “cone dystrophy with supernormal rod electroretinogram”. Invest Ophthalmol Vis Sci, 54, 898–908.

8. Zelinger, L., Wissinger, B., Eli, D., Kohl, S., Sharon, D. and Banin, E. (2013) Cone dystrophy with supernormal rod response: novel KCNV2 mutations in an underdiagnosed phenotype. Ophthalmology, 120, 2338–2343.

9. Czirjak, G., Toth, Z.E. and Enyedi, P. (2007) Characterization of the heteromeric potassium channel formed by kv2.1 and the retinal subunit kv8.2 in Xenopus oocytes. J Neurophysiol, 98, 1213–1222.

10. Gayet-Primo, J., Yaeger, D.B., Khanjian, R.A. and Puthussery, T. (2018) Heteromeric KV2/KV8.2 Channels Mediate Delayed Rectifier Potassium Currents in Primate Photoreceptors. J Neurosci, 38, 3414–3427.

11. Jiang, X., Rashwan, R., Voigt, V., Nerbonne, J., Hunt, D.M. and Carvalho, L.S. (2021) Molecular, Cellular and Functional Changes in the Retinas of Young Adult Mice Lacking the Voltage-Gated K(+) Channel Subunits Kv8.2 and K2.1. Int J Mol Sci, 22.

12. Hart, N.S., Mountford, J.K., Voigt, V., Fuller-Carter, P., Barth, M., Nerbonne, J.M., Hunt, D.M. and Carvalho, L.S. (2019) The Role of the Voltage-Gated Potassium Channel Proteins Kv8.2 and Kv2.1 in Vision and Retinal Disease: Insights from the Study of Mouse Gene Knock-Out Mutations. eNeuro, 6.

13. Fortenbach, C., Peinado Allina, G., Shores, C.M., Karlen, S.J., Miller, E.B., Bishop, H., Trimmer, J.S., Burns, M.E. and Pugh, E.N. (2021) Loss of the K+ channel Kv2.1 greatly reduces outward dark current and causes ionic dysregulation and degeneration in rod photoreceptors. J Gen Physiol, 153.

14. Smith, K.E., Wilkie, S.E., Tebbs-Warner, J.T., Jarvis, B.J., Gallasch, L., Stocker, M. and Hunt, D.M. (2012) Functional analysis of missense mutations in Kv8.2 causing cone dystrophy with supernormal rod electroretinogram. J Biol Chem, 287, 43972–43983.

15. McCulloch, D.L., Marmor, M.F., Brigell, M.G., Hamilton, R., Holder, G.E., Tzekov, R. and Bach, M. (2015) ISCEV Standard for full-field clinical electroretinography (2015 update). Doc Ophthalmol, 130, 1–12.

16. Joachimsthaler, A., Tsai, T.I. and Kremers, J. (2017) Electrophysiological Studies on The Dynamics of Luminance Adaptation in the Mouse Retina. Vision (Basel), 1.

17. Umino, Y., Guo, Y., Chen, C.K., Pasquale, R. and Solessio, E. (2019) Rod Photoresponse Kinetics Limit Temporal Contrast Sensitivity in Mesopic Vision. J Neurosci, 39, 3041–3056.

18. Volland, S., Esteve-Rudd, J., Hoo, J., Yee, C. and Williams, D.S. (2015) A comparison of some organizational characteristics of the mouse central retina and the human macula. PLoS One, 10, e0125631.

19. Kuburas, A., Thompson, S., Artemyev, N.O., Kardon, R.H. and Russo, A.F. (2014) Photophobia and abnormally sustained pupil responses in a mouse model of bradyopsia. Invest Ophthalmol Vis Sci, 55, 6878–6885.

20. Samardzija, M., Caprara, C., Heynen, S.R., Willcox DeParis, S., Meneau, I., Traber, G., Agca, C., von Lintig, J. and Grimm, C. (2014) A mouse model for studying cone photoreceptor pathologies. Invest Ophthalmol Vis Sci, 55, 5304–5313.

21. Barnes, S. (1994) After transduction: response shaping and control of transmission by ion channels of the photoreceptor inner segments. Neuroscience, 58, 447–459.

22. Yan, K. and Matthews, G. (1992) Blockers of potassium channels reduce the outward dark current in rod photoreceptor inner segments. Vis Neurosci, 8, 479–481.

23. Beech, D.J. and Barnes, S. (1989) Characterization of a voltage-gated K+ channel that accelerates the rod response to dim light. Neuron, 3, 573–581.

24. Grigg, J.R., Holder, G.E., Billson, F.A., Korsakova, M. and Jamieson, R.V. (2013) The importance of electrophysiology in revealing a complete homozygous deletion of KCNV2. J aapos, 17, 641–643.

25. Alexander, K.R. and Fishman, G.A. (1984) Supernormal scotopic ERG in cone dystrophy. Br J Ophthalmol, 68, 69–78.

26. Stockman, A., Henning, G.B., Michaelides, M., Moore, A.T., Webster, A.R., Cammack, J. and Ripamonti, C. (2014) Cone dystrophy with “supernormal” rod ERG: psychophysical testing shows comparable rod and cone temporal sensitivity losses with no gain in rod function. Invest Ophthalmol Vis Sci, 55, 832–840.

27. Witkovsky, P., Dudek, F.E. and Ripps, H. (1975) Slow PIII component of the carp electroretinogram. J Gen Physiol, 65, 119–134.

28. Lurie, M. and Marmor, M.F. (1980) Similarities between the c-wave and slow PIII in the rabbit eye. Invest Ophthalmol Vis Sci, 19, 1113–1117.

29. Lei, B. and Perlman, I. (1999) The contributions of voltage- and time-dependent potassium conductances to the electroretinogram in rabbits. Vis Neurosci, 16, 743–754.

30. Strettoi, E., Mears, A.J. and Swaroop, A. (2004) Recruitment of the rod pathway by cones in the absence of rods. J Neurosci, 24, 7576–7582.

31. Holter, P., Kunst, S., Wolloscheck, T., Kelleher, D.K., Sticht, C., Wolfrum, U. and Spessert, R. (2012) The retinal clock drives the expression of Kcnv2, a channel essential for visual function and cone survival. Invest Ophthalmol Vis Sci, 53, 6947–6954.

32. Jorge, B.S., Campbell, C.M., Miller, A.R., Rutter, E.D., Gurnett, C.A., Vanoye, C.G., George, A.L., Jr. and Kearney, J.A. (2011) Voltage-gated potassium channel KCNV2 (Kv8.2) contributes to epilepsy susceptibility. Proc Natl Acad Sci U S A, 108, 5443–5448.

33. Inamdar, S.M., Lankford, C.K., Laird, J.G., Novbatova, G., Tatro, N., Whitmore, S.S., Scheetz, T.E. and Baker, S.A. (2018) Analysis of 14-3-3 isoforms expressed in photoreceptors. Exp Eye Res, 170, 108–116.

34. Pan, Y., Bhattarai, S., Modestou, M., Drack, A.V., Chetkovich, D.M. and Baker, S.A. (2014) TRIP8b is required for maximal expression of HCN1 in the mouse retina. PLoS One, 9, e85850.

35. Vinberg, F. and Kefalov, V. (2015) Simultaneous ex vivo functional testing of two retinas by in vivo electroretinogram system. J Vis Exp, in press., e52855.

36. Kerov, V., Laird, J.G., Joiner, M.L., Knecht, S., Soh, D., Hagen, J., Gardner, S.H., Gutierrez, W., Yoshimatsu, T., Bhattarai, S. et al. (2018) alpha2delta-4 Is Required for the Molecular and Structural Organization of Rod and Cone Photoreceptor Synapses. J Neurosci, 38, 6145–6160.

