## Supplemental Statistical Tables for "Differential impact of Kv8.2 loss on rod and cone signaling and degeneration"

**Supplemental Statistical Tables** (Numbered according to corresponding Figure panel)

*Related to main Figures 3 and 9*

| **Table 3B: Sinusoidal Flicker ERG, magnitude of the fundamental** | | | | | | | | | | |
| --- | --- | --- | --- | --- | --- | --- | --- | --- | --- | --- |
| **Frequency (Hz)** | **WT** | | | **Kv8.2 KO** | | |  | **Sidak's multiple comparisons** | | |
|  | **Mean** | **SD** | **N** | **Mean** | **SD** | **N** | **Δ Mean** | **95% CI** | **Adj. P** | |
| 0.5 | 40.5 | 4.7 | 9 | 31.2 | 6.3 | 10 | 9.24 | 0.93,17.55 | 0.0231 | * |
| 0.75 | 37.4 | 5.8 | 9 | 26.5 | 5.8 | 10 | 10.9 | 2.17, 19.64 | 0.0089 | ** |
| 1 | 37.8 | 5.6 | 9 | 23.5 | 5.6 | 10 | 14.3 | 5.93, 22.72 | 0.0004 | *** |
| 1.5 | 36.2 | 5.4 | 9 | 18.5 | 5.0 | 10 | 17.8 | 9.90, 25.61 | <0.0001 | **** |
| 2 | 34.6 | 5.3 | 9 | 17.0 | 4.9 | 10 | 17.6 | 9.87, 25.28 | <0.0001 | **** |
| 3 | 28.6 | 5.1 | 9 | 13.9 | 3.6 | 10 | 14.6 | 7.82, 21.42 | <0.0001 | **** |
| 6 | 26.8 | 4.9 | 9 | 16.1 | 3.0 | 10 | 10.8 | 4.31, 17.18 | 0.0008 | *** |
| 9 | 34.3 | 6.2 | 9 | 24.3 | 4.9 | 10 | 10.0 | 1.45, 18.50 | 0.0158 | * |
| 12 | 27.0 | 6.5 | 9 | 11.0 | 3.0 | 10 | 15.9 | 7.57, 24.31 | 0.0004 | *** |
| 15 | 17.5 | 6.0 | 9 | 4.8 | 1.5 | 10 | 12.7 | 5.01, 20.47 | 0.0019 | ** |
| 25 | 3.1 | 1.2 | 9 | 0.7 | 0.2 | 9 | 2.4 | 0.89, 3.90 | 0.0028 | ** |

| **Table 3D: Sinusoidal Flicker ERG, b wave-like amplitude** | | | | | | | | | | |
| --- | --- | --- | --- | --- | --- | --- | --- | --- | --- | --- |
| **Frequency (Hz)** | **WT** | | | **Kv8.2 KO** | | |  | **Sidak's multiple comparisons** | | |
|  | **Mean** | **SD** | **N** | **Mean** | **SD** | **N** | **Δ Mean** | **95% CI** | **Adj. P** | |
| 0.5 | 181.8 | 18.1 | 9 | 270.8 | 33.7 | 10 | -89.0 | -122.1, -55.9 | <0.0001 | **** |
| 0.75 | 234.4 | 19.5 | 9 | 233.5 | 33.0 | 10 | 0.92 | -32.13, 33.96 | 0.9998 |  |
| 1 | 244.8 | 24.0 | 9 | 208.1 | 29.4 | 10 | 36.74 | 4.24, 69.23 | 0.0244 | * |

| **Table 3E: Sinusoidal Flicker ERG, c wave-like amplitude** | | | | | | | | | | |
| --- | --- | --- | --- | --- | --- | --- | --- | --- | --- | --- |
| **Frequency (Hz)** | **WT** | | | **Kv8.2 KO** | | |  | **Sidak's multiple comparisons** | | |
|  | **Mean** | **SD** | **N** | **Mean** | **SD** | **N** | **Δ Mean** | **95% CI** | **Adj. P** | |
| 0.5 | 197.4 | 30.4 | 9 | 93.1 | 14.3 | 10 | 104.40 | 73.27, 135.50 | <0.0001 | **** |
| 0.75 | 165.6 | 29.5 | 9 | 72.8 | 17.6 | 10 | 92.80 | 61.77, 123.80 | <0.0001 | **** |
| 1 | 132.8 | 24.1 | 9 | 72.8 | 11.6 | 10 | 60.00 | 35.24, 84.76 | <0.0001 | **** |

| **Table 9B: Photopic ERG a wave amplitude** | | | | | | | | | |
| --- | --- | --- | --- | --- | --- | --- | --- | --- | --- |
| **Stimulus**  **log(cd.s/m^2^)** | **Conefull: Kv8.2 WT** | | | **Conefull: Kv8.2 KO** | | |  | **Sidak's multiple comparisons** | |
|  | **Mean** | **SD** | **N** | **Mean** | **SD** | **N** | **Δ Mean** | **95% CI** | **Adj. P** |
| 0.0 | 4.1 | 3.6 | 6 | 3.5 | 3.1 | 13 | -0.61 | -6.74, 5.52 | >0.9999 |
| 0.40 | 2.5 | 1.1 | 6 | 4.1 | 3.3 | 13 | 1.70 | -1.60, 4.90 | 0.6562 |
| 0.9 | 4.3 | 1.7 | 6 | 4.2 | 3.6 | 13 | -0.07 | -3.80, 3.70 | >0.9999 |
| 1.40 | 4.6 | 2.0 | 6 | 6.4 | 2.6 | 13 | 1.89 | -1.72, 5.51 | 0.6114 |
| 1.90 | 3.7 | 1.3 | 6 | 5.3 | 3.1 | 13 | 1.62 | -1.49, 4.73 | 0.6498 |
| 2.40 | 7.0 | 3.9 | 6 | 12.7 | 6.9 | 13 | 5.66 | -2.13, 13.44 | 0.2593 |
| 2.90 | 20.9 | 9.2 | 6 | 17.9 | 6.2 | 13 | -3.00 | -18.67, 12.67 | 0.9954 |

| **Table 9C: Photopic ERG b wave amplitude** | | | | | | | | | | |
| --- | --- | --- | --- | --- | --- | --- | --- | --- | --- | --- |
| **Stimulus**  **log(cd.s/m^2^)** | **Conefull: Kv8.2 WT** | | | **Conefull: Kv8.2 KO** | | |  | **Sidak's multiple comparisons** | | |
|  | **Mean** | **SD** | **N** | **Mean** | **SD** | **N** | **Δ Mean** | **95% CI** | **Adj. P** | |
| 0.0 | 8.8 | 4.5 | 6 | 8.2 | 6.5 | 13 | -0.56 | -8.78, 7.67 | >0.9999 |  |
| 0.40 | 12.1 | 3.5 | 6 | 12.1 | 7.5 | 13 | 0.04 | -7.78, 7.87 | >0.9999 |  |
| 0.9 | 21.3 | 6.7 | 6 | 14.6 | 6.7 | 13 | -6.62 | -18.07, 4.83 | 0.4597 |  |
| 1.40 | 43.9 | 12.5 | 6 | 22.1 | 9.3 | 13 | -21.79 | -43.03, -0.55 | 0.0439 | * |
| 1.90 | 114.1 | 30.2 | 6 | 51.7 | 12.9 | 13 | -62.32 | -115.5, -9.11 | 0.0241 | * |
| 2.40 | 289.3 | 42.7 | 6 | 141.1 | 35.3 | 13 | -148.2 | -220.4, -75.93 | 0.0005 | *** |
| 2.90 | 449.0 | 35.1 | 6 | 257.5 | 57.2 | 13 | -191.5 | -259.1, -124.0 | <0.0001 | **** |

*Related to Supplemental Figures 4, 5, and 6*

| **Table S4B: Scotopic ERG a wave amplitude** | | | | | | | | | | |
| --- | --- | --- | --- | --- | --- | --- | --- | --- | --- | --- |
| **Stimulus**  **log(cd.s/m^2^)** | **WT** | | | **Kv8.2 KO** | | |  | **Sidak's multiple comparisons** | | |
|  | **Mean** | **SD** | **N** | **Mean** | **SD** | **N** | **Δ Mean** | **95% CI** | **Adj. P** | |
| -2.5 | 3.2 | 0.9 | 9 | 5.3 | 2.0 | 8 | -6.98 | -76.36, 62.40 | >0.9999 |  |
| -2.0 | 12.6 | 5.7 | 9 | 46.1 | 18.8 | 8 | -30.29 | -99.67, 39.09 | 0.8736 |  |
| -1.0 | 88.7 | 32.9 | 9 | 124.7 | 51.7 | 8 | -35.96 | -104, 32.12 | 0.7146 |  |
| 0.0 | 197.2 | 66.4 | 9 | 143.9 | 52.7 | 8 | 53.36 | -14.73, 121.4 | 0.2261 |  |
| 0.5 | 196.5 | 62.7 | 9 | 144.4 | 52.0 | 8 | 52.09 | -15.99, 120.2 | 0.2524 |  |
| 1.0 | 201.6 | 63.5 | 9 | 143.4 | 52.7 | 8 | 58.28 | -9.803, 126.4 | 0.1427 |  |
| 1.5 | 196.5 | 57.1 | 9 | 139.4 | 51.4 | 8 | 57.15 | -10.93, 125.2 | 0.1593 |  |
| 2.0 | 230.8 | 69.8 | 9 | 162.4 | 58.5 | 8 | 68.40 | 0.32, 136.5 | 0.0482 | * |

| **Table S4C: Scotopic ERG a wave implicit time** | | | | | | | | | | |
| --- | --- | --- | --- | --- | --- | --- | --- | --- | --- | --- |
| **Stimulus**  **log(cd.s/m^2^)** | **WT** | | | **Kv8.2 KO** | | |  | **Sidak's multiple comparisons** | | |
|  | **Mean** | **SD** | **N** | **Mean** | **SD** | **N** | **Δ Mean** | **95% CI** | **Adj. P** | |
| -2.5 | 30.9 | 2.7 | 9 | 73.3 | 10.3 | 8 | -42.38 | -56.25, -28.52 | <0.0001 | **** |
| -2.0 | 27.1 | 2.0 | 9 | 55.0 | 5.2 | 8 | -27.91 | -34.84, -20.99 | <0.0001 | **** |
| -1.0 | 21.8 | 1.4 | 9 | 36.2 | 2.6 | 8 | -14.39 | -17.91, -10.88 | <0.0001 | **** |
| 0.0 | 18.1 | 1.0 | 9 | 19.7 | 4.8 | 8 | -1.68 | -8.19, 4.83 | 0.9732 |  |
| 0.5 | 13.3 | 1.2 | 9 | 12.9 | 1.4 | 8 | 0.37 | -1.64, 2.37 | 0.9988 |  |
| 1.0 | 10.0 | 0.5 | 9 | 9.8 | 0.6 | 8 | 0.21 | -0.70, 1.13 | 0.9939 |  |
| 1.5 | 8.2 | 0.7 | 9 | 7.9 | 0.5 | 8 | 0.27 | -0.66, 1.19 | 0.9772 |  |
| 2.0 | 6.7 | 0.5 | 9 | 6.5 | 0.3 | 8 | 0.12 | -0.50, 0.73 | 0.9984 |  |

| **Table S4D: Scotopic ERG b wave amplitude** | | | | | | | | | |
| --- | --- | --- | --- | --- | --- | --- | --- | --- | --- |
| **Stimulus**  **log(cd.s/m^2^)** | **WT** | | | **Kv8.2 KO** | | |  | **Sidak's multiple comparisons** | |
|  | **Mean** | **SD** | **N** | **Mean** | **SD** | **N** | **Δ Mean** | **95% CI** | **Adj. P** |
| -2.5 | 90.3 | 17.8 | 9 | 163.4 | 23.6 | 8 | 36.59 | -118.1, 191.3 | 0.9968 |
| -2.0 | 76.0 | 13.0 | 9 | 151.2 | 9.3 | 8 | -74.20 | -228.9, 80.46 | 0.8063 |
| -1.0 | 44.6 | 5.8 | 9 | 86.8 | 7.0 | 8 | -78.19 | -232.9, 76.48 | 0.7594 |
| 0.0 | 40.9 | 3.0 | 9 | 66.2 | 5.9 | 8 | -13.70 | -168.4, 141.0 | >0.9999 |
| 0.5 | 41.6 | 2.2 | 9 | 61.2 | 5.2 | 8 | -8.47 | -163.1, 146.2 | >0.9999 |
| 1.0 | 43.1 | 2.1 | 9 | 58.2 | 4.7 | 8 | 3.60 | -151.1, 158.3 | >0.9999 |
| 1.5 | 44.7 | 1.2 | 9 | 56.3 | 4.9 | 8 | -6.17 | -160.8, 148.5 | >0.9999 |
| 2.0 | 39.7 | 5.3 | 9 | 57.6 | 3.3 | 8 | 42.27 | -112.4, 196.9 | 0.9916 |

| **Table S4E: Scotopic ERG b wave implicit time** | | | | | | | | | | |
| --- | --- | --- | --- | --- | --- | --- | --- | --- | --- | --- |
| **Stimulus**  **log(cd.s/m^2^)** | **WT** | | | **Kv8.2 KO** | | |  | **Sidak's multiple comparisons** | | |
|  | **Mean** | **SD** | **N** | **Mean** | **SD** | **N** | **Δ Mean** | **95% CI** | **Adj. P** | |
| -2.5 | 90.6 | 17.6 | 9 | 163.4 | 23.6 | 8 | -73.11 | -85.51, -60.70 | <0.0001 | **** |
| -2.0 | 76.0 | 13.0 | 9 | 151.2 | 9.3 | 8 | -75.24 | -87.64, -62.83 | <0.0001 | **** |
| -1.0 | 44.6 | 5.8 | 9 | 86.8 | 7.0 | 8 | -42.19 | -54.60, -29.79 | <0.0001 | **** |
| 0.0 | 40.9 | 3.0 | 9 | 66.2 | 5.3 | 8 | -25.27 | -37.67, -12.86 | <0.0001 | **** |
| 0.5 | 41.6 | 2.2 | 9 | 61.2 | 5.2 | 8 | -19.55 | -31.96, -7.15 | 0.0002 | *** |
| 1.0 | 43.1 | 2.1 | 9 | 58.2 | 4.7 | 8 | -15.16 | -27.56, -2.76 | 0.0075 | ** |
| 1.5 | 44.7 | 2.0 | 9 | 56.3 | 4.9 | 8 | -11.65 | -24.05, 0.76 | 0.0797 |  |
| 2.0 | 39.7 | 5.3 | 9 | 57.6 | 3.3 | 8 | -17.88 | -30.29, -5.48 | 0.0009 | *** |

| **Table S4F: Scotopic ERG b/a wave** | | | | | | | | | | |
| --- | --- | --- | --- | --- | --- | --- | --- | --- | --- | --- |
| **Stimulus**  **log(cd.s/m^2^)** | **WT** | | | **Kv8.2 KO** | | |  | **Sidak's multiple comparisons** | | |
|  | **Mean** | **SD** | **N** | **Mean** | **SD** | **N** | **Δ Mean** | **95% CI** | **Adj. P** | |
| -2.5 | 66.4 | 35.7 | 9 | 28.7 | 9.5 | 8 | 37.75 | -5.58, 81.09 | 0.1017 |  |
| -2.0 | 24.1 | 7.8 | 9 | 8.5 | 1.5 | 8 | 15.50 | 4.50, 26.11 | 0.0056 | ** |
| -1.0 | 3.5 | 7.8 | 9 | 8.5 | 1.5 | 8 | 0.38 | -0.50, 1.26 | 0.7944 |  |
| 0.0 | 2.0 | 0.2 | 9 | 2.8 | 0.2 | 8 | -0.08 | -1.15, -0.05 | <0.0001 | **** |
| 0.5 | 2.1 | 0.1 | 9 | 2.9 | 0.2 | 8 | -0.80 | -1.05, -0.55 | <0.0001 | **** |
| 1.0 | 2.2 | 0.2 | 9 | 3.0 | 0.2 | 8 | -0.87 | -1.18, -0.56 | <0.0001 | **** |
| 1.5 | 2.1 | 0.2 | 9 | 3.1 | 0.3 | 8 | -0.97 | -1.33, -0.60 | <0.0001 | **** |
| 2.0 | 2.2 | 0.2 | 9 | 2.9 | 0.2 | 8 | -0.69 | -1.04, -0.34 | 0.0002 | *** |

| **Table S4G: Scotopic ERG a minus b wave implicit time** | | | | | | | | | | |
| --- | --- | --- | --- | --- | --- | --- | --- | --- | --- | --- |
| **Stimulus**  **log(cd.s/m^2^)** | **WT** | | | **Kv8.2 KO** | | |  | **Sidak's multiple comparisons** | | |
|  | **Mean** | **SD** | **N** | **Mean** | **SD** | **N** | **Δ Mean** | **95% CI** | **Adj. P** | |
| -2.5 | 59 | 16 | 9 | 90 | 16 | 8 | -30.73 | -55.47, -5.98 | 0.0107 | * |
| -2.0 | 49 | 12 | 9 | 96 | 5 | 8 | -47.36 | -61.75, -32.97 | <0.0001 | **** |
| -1.0 | 23 | 5 | 9 | 51 | 7 | 8 | -27.80 | -37.40, -18.20 | <0.0001 | **** |
| 0.0 | 23 | 2 | 9 | 46 | 6 | 8 | -23.57 | -32.16, -14.99 | <0.0001 | **** |
| 0.5 | 28 | 2 | 9 | 48 | 6 | 8 | -19.88 | -27.30, -12.45 | <0.0001 | **** |
| 1.0 | 33 | 2 | 9 | 48 | 5 | 8 | -15.41 | -21.62, -9.20 | <0.0001 | **** |
| 1.5 | 36 | 2 | 9 | 48 | 5 | 8 | -11.94 | -18.29, -5.58 | 0.0008 | *** |
| 2.0 | 33 | 5 | 9 | 51 | 3 | 8 | -17.99 | -24.48, -11.50 | <0.0001 | **** |

| **Table S5B: Photopic ERG a wave amplitude** | | | | | | | | | | |
| --- | --- | --- | --- | --- | --- | --- | --- | --- | --- | --- |
| **Stimulus**  **log(cd.s/m^2^)** | **WT** | | | **Kv8.2 KO** | | |  | **Sidak's multiple comparisons** | | |
|  | **Mean** | **SD** | **N** | **Mean** | **SD** | **N** | **Δ Mean** | **95% CI** | **Adj. P** | |
| 0.0 | 5.0 | 3.5 | 9 | 4.0 | 1.9 | 8 | 1.30 | -2.98, 5.56 | 0.9530 |  |
| 0.40 | 9.0 | 4.0 | 9 | 8.3 | 4.5 | 8 | 0.68 | -5.79, 7.15 | >0.9999 |  |
| 0.9 | 14.1 | 3.8 | 9 | 9.7 | 5.5 | 8 | 4.40 | -3.04, 11.83 | 0.448 |  |
| 1.40 | 18.2 | 7.0 | 9 | 10.6 | 2.6 | 8 | 7.57 | -0.49, 15.63 | 0.0703 |  |
| 1.90 | 20.8 | 6.1 | 9 | 11.4 | 2.2 | 8 | 9.34 | 2.08, 16.59 | 0.0103 | * |
| 2.40 | 23.0 | 5.3 | 9 | 15.0 | 3.6 | 8 | 7.98 | 1.14, 14.83 | 0.0173 | * |
| 2.90 | 29.2 | 7.2 | 9 | 19.7 | 5.00 | 8 | 9.51 | 0.20, 18.82 | 0.0437 | * |

| **Table S5C: Photopic ERG a wave implicit time** | | | | | | | | | | |
| --- | --- | --- | --- | --- | --- | --- | --- | --- | --- | --- |
| **Stimulus**  **log(cd.s/m^2^)** | **WT** | | | **Kv8.2 KO** | | |  | **Sidak's multiple comparisons** | | |
|  | **Mean** | **SD** | **N** | **Mean** | **SD** | **N** | **Δ Mean** | **95% CI** | **Adj. P** | |
| 0.0 | 16.4 | 5.2 | 9 | 22.1 | 6.7 | 8 | -5.74 | -15.01, 3.53 | 0.4052 |  |
| 0.40 | 17.5 | 1.3 | 9 | 22.2 | 2.2 | 8 | -4.75 | -7.7, -1.80 | 0.0017 | ** |
| 0.9 | 15.9 | 1.3 | 9 | 19.1 | 4.9 | 8 | -3.18 | -9.58, 3.23 | 0.5712 |  |
| 1.40 | 14.0 | 0.9 | 9 | 17.6 | 2 | 8 | -3.55 | 6.17, 0.94 | 0.0075 | ** |
| 1.90 | 12.3 | 0.9 | 9 | 15.3 | 3.0 | 8 | -3.02 | -6.95, 0.92 | 0.1675 |  |
| 2.40 | 11.1 | 0.5 | 9 | 14.1 | 3.7 | 8 | -2.95 | -7.78, 1.88 | 0.3397 |  |
| 2.90 | 9.7 | 1.3 | 9 | 10.9 | 3.4 | 8 | 1.26 | -5.72, 3.21 | 0.9540 |  |

| **Table S5D: Photopic ERG b wave amplitude** | | | | | | | | | | |
| --- | --- | --- | --- | --- | --- | --- | --- | --- | --- | --- |
| **Stimulus**  **log(cd.s/m^2^)** | **WT** | | | **Kv8.2 KO** | | |  | **Sidak's multiple comparisons** | | |
|  | **Mean** | **SD** | **N** | **Mean** | **SD** | **N** | **Δ Mean** | **95% CI** | **Adj. P** | |
| 0.0 | 35.4 | 12.6 | 9 | 13.2 | 3.7 | 8 | 22.29 | 7.31, 37.27 | 0.0040 | ** |
| 0.40 | 72.1 | 22.4 | 9 | 24.8 | 8.9 | 8 | 47.33 | 20.54, 74.11 | 0.0009 | *** |
| 0.9 | 138.3 | 36.9 | 9 | 46.6 | 15.6 | 8 | 91.73 | 47.48, 136 | 0.0002 | *** |
| 1.40 | 150.8 | 37.7 | 9 | 57.6 | 16.4 | 8 | 93.25 | 47.99, 138.5 | 0.0002 | *** |
| 1.90 | 155.5 | 41.5 | 9 | 62.2 | 17.5 | 8 | 93.31 | 43.60, 143 | 0.0005 | *** |
| 2.40 | 168.5 | 43.3 | 9 | 67.7 | 17.5 | 8 | 100.9 | 49.07, 152.6 | 0.0004 | *** |
| 2.90 | 186.1 | 47.2 | 9 | 77.4 | 20.2 | 8 | 108.7 | 52.14, 165.3 | 0.0004 | *** |

| **Table S5E: Photopic ERG b wave implicit time** | | | | | | | | | | |
| --- | --- | --- | --- | --- | --- | --- | --- | --- | --- | --- |
| **Stimulus**  **log(cd.s/m^2^)** | **WT** | | | **Kv8.2 KO** | | |  | **Sidak's multiple comparisons** | | |
|  | **Mean** | **SD** | **N** | **Mean** | **SD** | **N** | **Δ Mean** | **95% CI** | **Adj. P** | |
| 0.0 | 34.9 | 1.7 | 9 | 41.5 | 4.6 | 8 | -6.63 | -12.59, 0.66 | 0.0281 | * |
| 0.40 | 43.4 | 3.6 | 9 | 46.8 | 5.6 | 8 | -3.35 | -10.83, 4.13 | 0.7354 |  |
| 0.9 | 41.5 | 2.2 | 9 | 47.0 | 5.8 | 8 | -5.47 | -13.01, 2.07 | 0.2109 |  |
| 1.40 | 39.5 | 2.1 | 9 | 48.9 | 5.4 | 8 | -9.39 | -16.45, -2.33 | 0.0093 | ** |
| 1.90 | 38 | 2.0 | 9 | 46.9 | 5.1 | 8 | -8.89 | -15.58, -2.20 | 0.0093 | ** |
| 2.40 | 36.5 | 1.8 | 9 | 45.5 | 5.2 | 8 | -8.96 | -15.74, -2.17 | 0.0101 | * |
| 2.90 | 34.8 | 1.7 | 9 | 43.7 | 4.9 | 8 | -8.90 | -15.31, -2.50 | 0.0074 | ** |

| **Table S5F: Photopic ERG b/a wave** | | | | | | | | | |
| --- | --- | --- | --- | --- | --- | --- | --- | --- | --- |
| **Stimulus**  **log(cd.s/m^2^)** | **WT** | | | **Kv8.2 KO** | | |  | **Sidak's multiple comparisons** | |
|  | **Mean** | **SD** | **N** | **Mean** | **SD** | **N** | **Δ Mean** | **95% CI** | **Adj. P** |
| 0.0 | 14.1 | 15.5 | 9 | 3.5 | 1.9 | 7 | 10.60 | -7.75, 28.96 | 0.4192 |
| 0.40 | 10.0 | 5.9 | 9 | 4.2 | 3.0 | 8 | 5.81 | -1.37, 13 | 0.1498 |
| 0.9 | 9.8 | 0.8 | 9 | 6.5 | 6.1 | 7 | 3.33 | -5.77, 12.43 | 0.7890 |
| 1.40 | 8.6 | 1.6 | 9 | 5.8 | 2.7 | 8 | 2.81 | -0.73, 6.34 | 0.1594 |
| 1.90 | 7.6 | 0.7 | 9 | 5.8 | 2.5 | 8 | 1.78 | -1.51, 5.06 | 0.4787 |
| 2.40 | 7.4 | 0.8 | 9 | 5.0 | 2.4 | 8 | 2.33 | -0.83, 5.49 | 0.1977 |
| 2.90 | 6.4 | 0.7 | 9 | 4.3 | 1.8 | 8 | 2.15 | -0.17, 4.47 | 0.0739 |

| **Table S5G: Photopic ERG a minus b wave implicit time** | | | | | | | | | | |
| --- | --- | --- | --- | --- | --- | --- | --- | --- | --- | --- |
| **Stimulus**  **log(cd.s/m^2^)** | **WT** | | | **Kv8.2 KO** | | |  | **Sidak's multiple comparisons** | | |
|  | **Mean** | **SD** | **N** | **Mean** | **SD** | **N** | **Δ Mean** | **95% CI** | **Adj. P** | |
| 0.0 | 18.5 | 6.0 | 9 | 19.4 | 7.3 | 8 | 0.89 | -11.20, 9.42 | >0.9999 |  |
| 0.40 | 25.9 | 3.0 | 9 | 24.5 | 6.1 | 8 | 1.4 | -6.56, 9.36 | 0.9972 |  |
| 0.9 | 25.6 | 1.4 | 9 | 27.9 | 10.0 | 8 | -2.3 | -15.39, 10.80 | 0.9954 |  |
| 1.40 | 25.5 | 1.5 | 9 | 31.3 | 5.3 | 8 | -5.8 | -12.73, 1.06 | 0.1103 |  |
| 1.90 | 25.7 | 1.3 | 9 | 31.6 | 5.7 | 8 | -5.9 | -13.29, 1.54 | 0.1438 |  |
| 2.40 | 25.4 | 1.5 | 9 | 31.5 | 6.0 | 8 | -6.01 | 13.86, 1.84 | 0.1651 |  |
| 2.90 | 25.2 | 1.6 | 9 | 32.8 | 4.7 | 8 | -7.65 | 13.8, 1.50 | 0.0147 | * |

| **Table S6: Flash Flicker ERG** | | | | | | | | | | |
| --- | --- | --- | --- | --- | --- | --- | --- | --- | --- | --- |
| **Frequency (Hz)** | **WT** | | | **Kv8.2 KO** | | |  | **Sidak's multiple comparisons** | | |
|  | **Mean** | **SD** | **N** | **Mean** | **SD** | **N** | **Δ Mean** | **95% CI** | **Adj. P** | |
| 0.5 | 471.6 | 107.0 | 7 | 453.4 | 60.2 | 8 | 18.23 | -154.1, 190.5 | >0.9999 |  |
| 0.75 | 345.6 | 93.7 | 7 | 216.9 | 29.7 | 8 | 128.80 | -24.15, 281.7 | 0.1140 |  |
| 1 | 206.6 | 51.4 | 7 | 103.8 | 9.5 | 8 | 102.80 | 17.22, 188.5 | 0.0198 | * |
| 1.5 | 169.4 | 37.6 | 7 | 96.0 | 9.0 | 8 | 73.39 | 11.35, 135.4 | 0.0211 | * |
| 2 | 175.2 | 28.3 | 7 | 96.0 | 16.1 | 8 | 79.16 | 33.56, 124.8 | 0.0011 | ** |
| 3 | 159.4 | 24.9 | 7 | 83.2 | 12.1 | 8 | 76.23 | 36.19, 116.3 | 0.0007 | *** |
| 6 | 124.5 | 31.4 | 7 | 62.3 | 12.7 | 8 | 62.13 | 11.38, 112.9 | 0.0159 | * |
| 9 | 126.9 | 29.9 | 7 | 61.2 | 15.5 | 8 | 65.76 | 17.73, 113.8 | 0.0070 | ** |
| 12 | 103.1 | 24.1 | 7 | 39.9 | 9.1 | 8 | 63.16 | 24.02, 102.3 | 0.0029 | ** |
| 15 | 77.1 | 16.1 | 7 | 35.7 | 6.1 | 8 | 41.37 | 15.22, 67.52 | 0.0033 | ** |
| 25 | 64.0 | 11.3 | 7 | 30.9 | 5.8 | 8 | 33.04 | 14.87, 51.22 | 0.0009 | *** |
| 30 | 48.4 | 10.5 | 7 | 13.2 | 3.2 | 8 | 35.20 | 18.06, 52.34 | 0.0007 | *** |
